# Netrins and receptors control *Drosophila* optic lobe organization and transmedullary neuron axon targeting

**DOI:** 10.1101/2021.10.30.466625

**Authors:** Yu Zhang, Scott Lowe, Xin Li

## Abstract

During development, integration of temporal patterning and spatial patterning of neural progenitors as well as Notch-dependent binary fate choice between sister neurons contribute to generation of neural diversity. How these upstream neural fate specification programs regulate downstream effector genes to control axon targeting and neuropil assembly remains less well-understood. Here we show that Notch-dependent binary fate choice in *Drosophila* medulla neurons regulates the expression of Netrin, and that Netrin pathway controls axon guidance of transmedullary (Tm) neurons and contributes to the organization of optic lobe neuropils. Netrins are enriched in the lobula where Tm axons target, and the attractive receptor Frazzled is expressed broadly in medulla neurons, while the repulsive receptor Unc-5 is excluded from Tm neurons and this is necessary for their correct targeting to the lobula. Frazzled is required collectively in a group of early-born Tm neurons to establish the inner optic chiasm (IOC) through which Tm axons target lobula. In addition, Frazzled acts in the layer-specific targeting step of Tm3 and Tm4 cell-autonomously, and is also required for the formation of the lobula branch of TmY3. Moreover, we show that the diffusibility of Netrins is necessary for Netrin enrichment in the lobula, the IOC formation and layer-specific targeting of Tm3 and Tm4. Netrin enrichment in the lobula is promoted by Frazzled expressed in Tm neurons, while Unc-5 appears to have an opposite role in Netrin distribution. Furthermore, we show that Netrin B is expressed in the Notch-on hemilineage of medulla neurons including most Tm and TmY neurons that target lobula, and loss of Su(H) abolished NetB expression in the medulla. Without medulla-originated NetB, Tm axons from late-born medulla columns cannot join the IOC. Therefore, the Notch-dependent binary fate choice regulates the assembly of the optic lobe neuropils by controlling the expression of Netrin.

## Introduction

During neural development, a great diversity of neural types are specified and their axons navigate for a long distance to connect to their correct targets, which are critical steps in the development of neural circuits and assembly of neuropils. A large number of studies have focused on how neural diversity generation is controlled by temporal and spatial patterning of neural progenitors and progeny by gene regulatory networks and signaling pathways [reviewed in ^1–7^]. In parallel, a wealth of studies over the past several decades have elaborated on the cellular machinery that shape the neurons: extracellular guidance cues, such as Netrins, Slits, Semaphorins, Ephrins, certain morphogens and growth factors, as well as cell-adhesion molecules, act on their receptors or other cell-adhesion molecules on the cell surface; these cell-surface molecules are linked to intracellular signaling proteins, which then regulate the cytoskeleton dynamics [reviewed in ^8–10^]. For example, Netrin-1 (NetA and NetB in *Drosophila*) is a conserved guidance cue, and Netrin binding induces homodimerization of DCC receptor (Frazzled in *Drosophila*), which leads to recruitment of downstream effectors and regulatory proteins including Trim9, and finally causes attractive response. In contrast, heterodimerization between DCC/Fra and receptor uncoordinated locomotion 5 (UNC5) leads to repulsion [reviewed in ^11^]. However, studies linking the two fields are relatively lacking, and it is not well-understood how the upstream neural fate specification programs are “translated into” the highly-organized neuropils by controlling coordinated axon targeting through regulating downstream effectors such as guidance cues and cell surface molecules in different groups of neurons.

We address this question using the *Drosophila* visual system, which has been an excellent model system to study the mechanism of axon guidance. The adult *Drosophila* visual system is composed of the compound eye with ~800 ommatidia containing inner and outer photoreceptors (PRs), and the optic lobe, which processes visual information and relays it to the central brain. The optic lobe consists of four neuropils, lamina, medulla, lobula and lobula plate, and each neuropil is comprised of ~800 units (columns or cartridges). Lamina neurons receive inputs from outer PRs; axons of lamina neurons and inner PRs project to the medulla neuropil through the outer optic chiasm (ooc). The medulla, lobula and lobula plate neuropils can be divided into 10 (M1-M10), 6 (lo1-lo6), and 4 (lop1-lop4) layers, respectively. As the largest neuropil in the optic lobe, the medulla consists of ~40,000 neurons classified into more than 80 neural types, belonging to several broad classes including Medulla intrinsic neurons (Mi), Distal medulla neurons (Dm), Proximal medulla neurons (Pm), Transmedullary neurons (Tm, connecting medulla with lobula), TmY (connecting medulla with lobula and lobula plate), Mt neurons and others ^12–14^. During development, medulla neurons are generated by medulla neuroblasts sequentially transformed from the neuroepithelial cells (NE) at the medial Outer Proliferation Center (OPC). Medulla neuron diversity is achieved mainly by the integration of the spatial and temporal patterning of medulla neuroblasts and Notch-dependent binary fate choices. Temporal patterning is mediated by a cascade of temporal transcription factors (TTF), including Homothorax (Hth), Eyeless (Ey), Sloppy paired 1 and 2 (Slp), Dichaete (D) and Tailless (Tll) in neuroblasts ^15,16^. More TTFs have been found by scRNA-seq of medulla neuroblasts recently ^17–19^. Notch-dependent binary fate choice happens in two daughters of GMCs. Each GMC gives rise to two daughters of distinct fates, with one daughter turning on the Notch (N) signaling and subsequently the transcription factor, Apterous (Ap), and the other daughter without N signaling^15^. Among medulla neurons, Tm and TmY neurons project their axons through the Inner optic chiasm (IOC) to the Lobula and lobula plate, which then send visual information to the central brain. Due to the progressive neurogenesis wave in the OPC, there is a temporal gradient of Tm neuron targeting and lobula column assembly, with Tms from the most anterior medulla columns target to the most proximal lobula columns first, followed by later-born Tms from more posterior medulla columns targeting to distal lobula columns, thus forming the IOC ^20^ (Fig. 1a). EM reconstruction of IOC revealed interleaving bundles and sheets of axons, with axon bundles each containing Tm axons from multiple columns connecting medulla and the lobula complex, and axon sheets connecting lobula and lobula plate ^21^. When Tm axons from posterior medulla columns leave the bundle, they make a turn to join the sheets of axons, and pass through anterior bundles forming the IOC, to project to the lobula ^21^.

**Figure 1.**
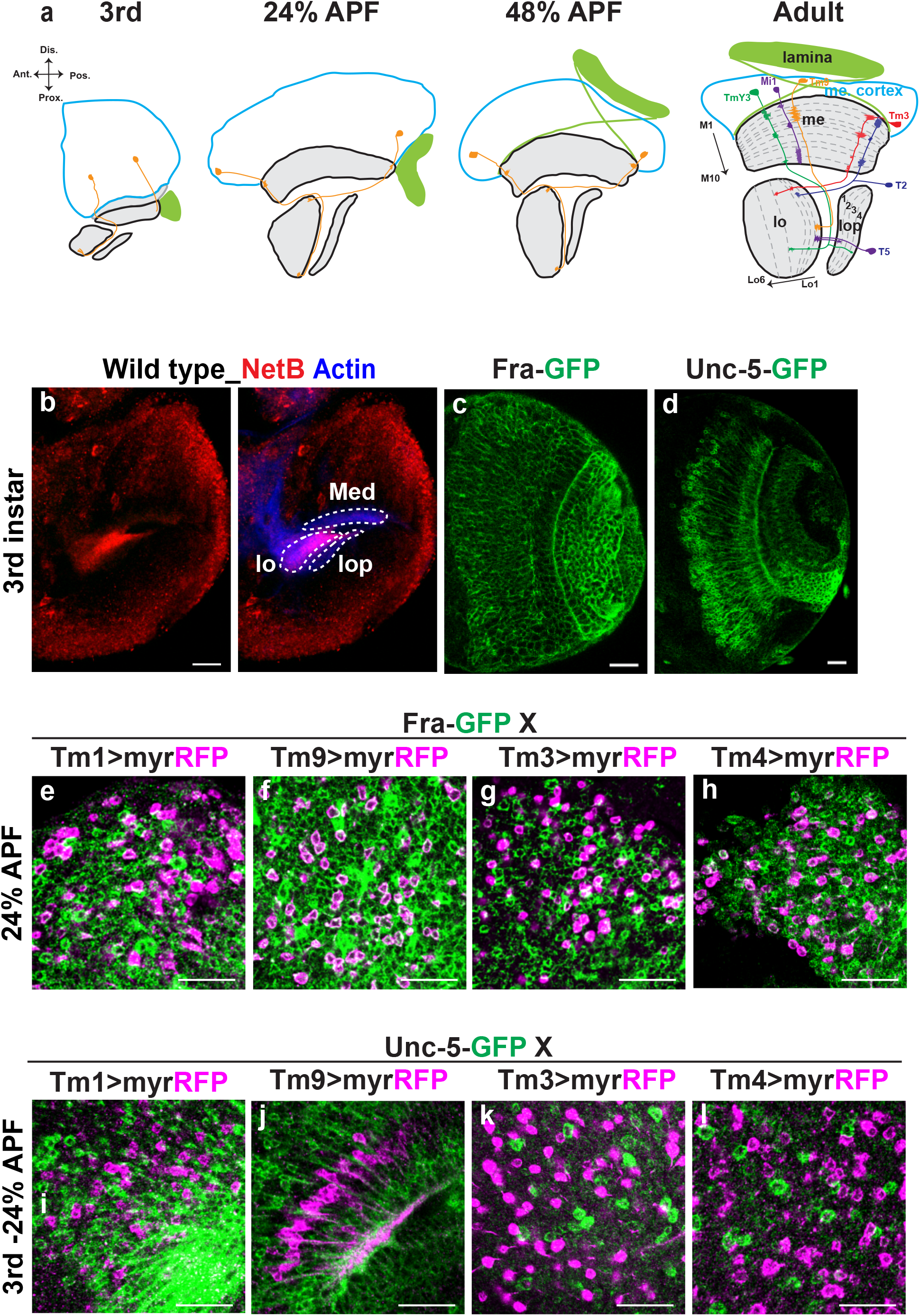
Netrins, Fra and Unc-5 are expressed in the optic lobe during development. (a) A schematic drawing of the optic lobe from the 3^rd^ instar larval stage to adult. Me: medulla. Lo: lobula. Lop: lobula plate. (b) The expression/distribution pattern of NetB (red) in the 3^rd^ larval optic lobe. Actin is labeled by phalloidin (blue). Neuropils are outlined. (c) The expression pattern of Fra∷GFP (green) in 3^rd^ larval optic lobe. (d) The expression pattern of Unc-5∷GFP (green) in 3^rd^ larval optic lobe. (e-h) 24% APF pupa brains. Tm1, Tm9, Tm3 and Tm4 neurons (purple) express Fra∷GFP (Green). (i-l) 3^rd^ to 24% APF brains. Tm1, Tm9, Tm3 and Tm4 (purple) neurons do not express Unc-5∷GFP (Green). Scale bar: 20μm.

Studies of axon guidance in the *Drosophila* visual system used to focus on photoreceptors and lamina neurons. These studies revealed a large number of cell surface molecules and transcription factors involved in the guidance of axons, formation of columns and layers, and synaptic specificity ^22–24^. For one example, transcription factor dFezf /Erm is required for L3 to target M3 layer, and also for its expression of Netrin, which is captured and concentrated at layer M3 ^25,26^. Netrin receptor Fra is then required in R8 axons to stably attach to Netrin that is concentrated at the M3 layer ^25,27^. Recently, several scRNA-seq studies of pupal and adult optic lobes have successfully matched developmental transcriptome profiles of more than 60 cell types to known morphological cell types^28–32^. Among the most differentially expressed genes, transcription factors and cell surface proteins are highly informative, providing a valuable resource to decode the mechanism of axon guidance. For a specific cell type, which cell surface proteins are required for axon guidance and how specific transcription factors regulate them are the next questions to answer. For example, in the eight subtypes of T4/T5 neurons, modular transcription factor codes were found to separately define axon and dendrite connectivity possibly by regulating the expression of combinations of cell surface proteins^32^.

However, for medulla neurons, especially Tms, cell surface molecules involved in axon guidance and how they are regulated remain largely unknown. Moreover, how the global organization of neuropils and the formation of IOC is regulated by upstream neuron specification programs remains unclear.

Here, we report that Netrins are required for the axon guidance of Tm neurons to the lobula and for the IOC formation. Fra is required collectively in a group of pioneering Tm axons born in early temporal stages to establish the IOC, and is also required cell-autonomously for layer specificity in a subset of Tm neurons that target to deep lobula layers. Fra expressed in Tm neurons is collectively required to enrich NetB in the lobula, while the presence of Unc-5 negatively affects enrichment of NetB in neuropils. Interestingly, the expression of NetB is activated by Notch signaling in medulla neurons of N-on hemi-lineage composed of mostly Tm/TmY neurons that target the lobula. NetB provided by N-on medulla neurons is required to facilitate the axon guidance of Tms from later-generated columns. This study identifies a mechanism for connecting upstream neuron specification programs to downstream effector genes and the global organization of neuropils.

## Results

### Netrins and receptors are expressed in the optic lobe during development

From the early third-instar larval stage, medulla neuroblasts divide to generate most medulla neurons. Medulla neurons start to extend their axons shortly after birth, and at around 55% after puparium formation (APF), most of them arrive at their target layers, ready to form synapses and showing the stimulus-independent neural activity^33^. Among them, axons of transmedullary neurons extend through the IOC to target the lobula^20^ (Fig. 1a). To identify the guidance cue required for Tm neuron targeting, we examined the expression patterns of known guidance cues, and found that Netrin-B (NetB) is exclusively enriched in the lobula (Fig. 1b,b’), while Netrin-A (NetA) is enriched at the boundaries around neuropils (Supplementary Fig. 1a). Then we examined the expression pattern of Netrin receptors in the third-instar larval medulla cortex, where cell bodies of medulla neurons reside. We found that the two Netrin receptors, Frazzled (Fra) and Unc-5, were expressed in the medulla cortex at the third-instar larval stage. Fra is broadly expressed in the medulla cortex (Fig. 1c), while Unc-5 is expressed in subset of cells (Fig. 1d). The GFP-tagged line and antibody staining of Fra were confirmed to show the same expression patterns (Supplementary Fig. 1 b, c).

Previous studies found that cells expressing Fra but not Unc-5 are attracted to the Netrin source, while Unc-5-expressing cells are repelled from the Netrin source^34,35^. Since NetB is enriched in the lobula neuropil at the beginning of the targeting stage, we speculated that medulla neurons that project to the lobula neuropil, i.e., transmedullary neurons (Tm), should express the attractive receptor Fra, but not the repulsive receptor Unc-5. We examined Fra expression in Tm neurons at 24% APF when most of them are extending their axons to the lobula. Using cell-type-specific drivers, we found that Tm1^13^, Tm3, Tm4^30^, and Tm9^30,36^ express Fra at 24% APF (Fig. 1e-h). Consistent with our hypothesis, none of the Tm neurons express the repulsive Unc-5 receptor (Fig. 1i-l). We identified a Tm3-specific gal4 driver (GMR12C11-GAL4) that starts expression at late 3^rd^ larval stage, and the identity of labeled neurons was confirmed by staining with Tm3 identity markers (Dfr and Cut)^37^ at the early pupal stage and by co-staining with existing Tm3-lexA^38^ at the adult stage (Supplementary Fig. 1d, e). Moreover, using molecular markers, we found Tm5a/b (Ap+, Toy+, and Sox102F+)^28,39^ do not express Unc-5 (Supplementary Fig. 1f). In summary, Netrins are enriched in the lobula neuropil during Tm axon targeting, and axons of various Tm neurons which target the lobula neuropil express the attractive receptor Fra but not the repulsive receptor Unc-5.

### Fra is required cell-autonomously for Tm3 and Tm4 targeting to deep lobula layers

Since Fra was expressed in Tm neurons that target the Netrin concentrated lobula neuropil, we next checked whether Fra was required to target Tm axons. By crossing UAS-fraRNAi with cell-type-specific drivers, we knocked down fra specifically in different Tm neurons. We found that targeting of Tm1 and Tm9 to lo1 was not affected by loss of fra (Supplementary Fig. 2b, e). In contrast, 65.4% of Tm3 and 72% of Tm4 axons were mistargeted to lo2 or lobula plate (lop) comparing with all Tm3 and Tm4 axons targeting to lo4 in wild type (Fig. 2b, h). The knockdown efficiency of UAS-fra RNAi was confirmed by anti-Fra staining (Supplementary Fig. 2h).

**Figure 2.**
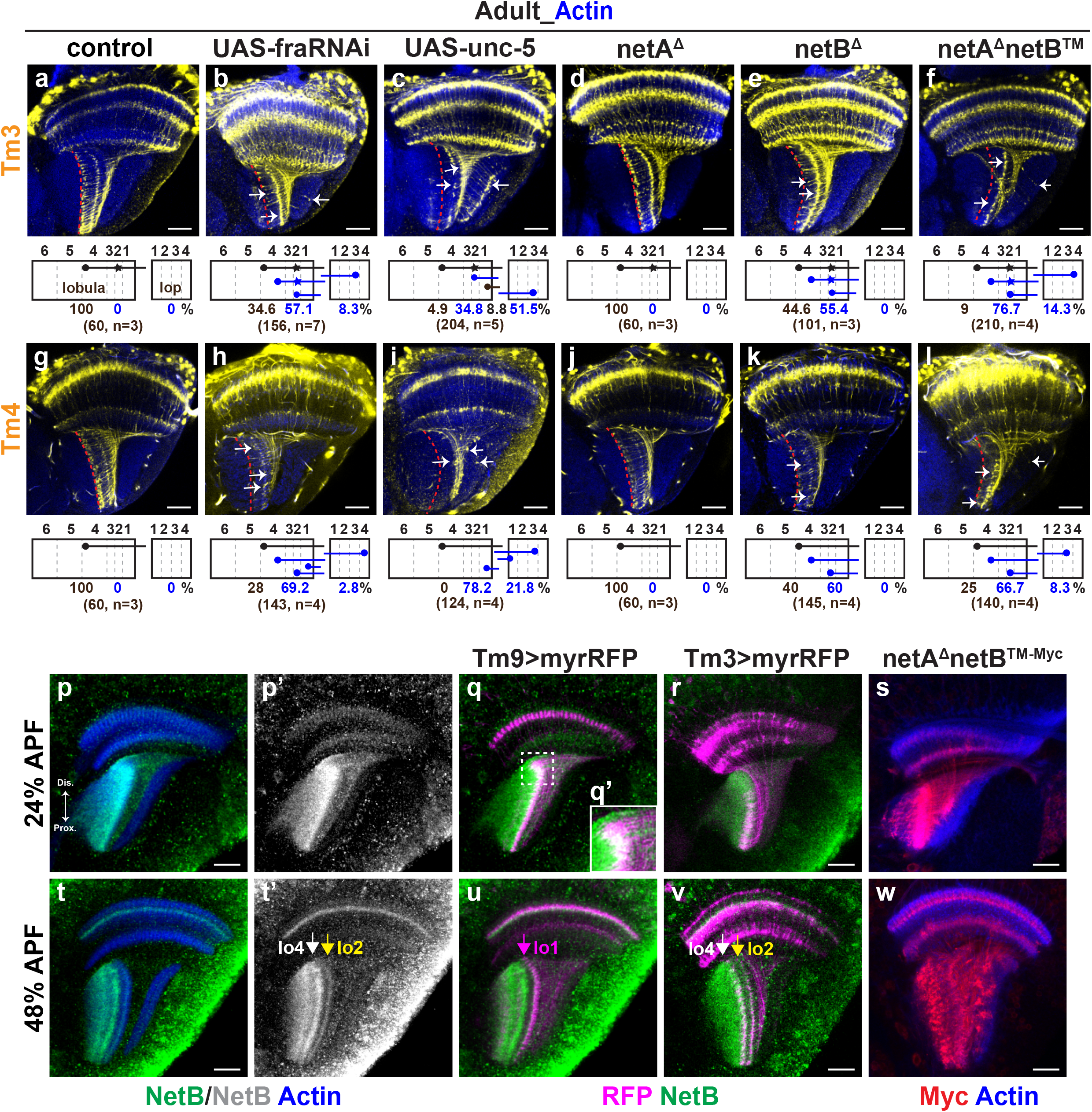
Netrin signaling is required for layer-specific targeting of Tm3 and Tm4. (a-f) The red dashed line marks the wild-type Tm3 axon targeting layer. Arrows point to the mistargeted axon terminals. (a-c) Tm3 are labeled by early gal4 (BL76324) with *UAS-myrRFP*. (d-f) Tm3 are labeled by *Tm3-lexA>lexop-tdTomato*. Schematic below each image shows the six layered lobula (left, lo1-lo6), four layered lobula plate (right, lop.1-lop.4) and Tm3 arborization (axon terminals are shown as black dots, and arbors at lo2 are shown as star). Percentage of targeting layers, total number of counted axons and the sample size are also shown. (a) Wild-type Tm3 axons (yellow) arborize at lo2 and terminate at lo4 (red dashed line) (b) Knocking down Fra specifically in Tm3. Lo4 layer labeled by Tm3 is not continuous, indicating that not all Tm3 terminate at lo4. (c) Overexpressing Unc-5 specifically in Tm3. Most Tm3 mistargeted at lop, lo1-lo2. (d) Tm3 targeting in NetA mutants is normal. (e) Tm3 targeting in NetB mutants is disrupted. (f) Tm3 targeting in indiffusible NetB mutants is abnormal. Most Tm3 stall at lo2-lo3. (g-l) Tm4 are labelled by type-specific gal4 with *uas-mcd8GFP*. Schematic and quantification are shown below. The red dashed line labels the wild-type Tm4 axon targeting layer. (g) Wild-type Tm4 (yellow) terminate at lo4 (red dashed line). (h) Knocking down Fra specifically in Tm4. Lo4 layer labeled by Tm4 is not continuous, indicating that not all Tm4 terminate at lo4. (i) Overexpressing Unc-5 specifically in Tm3. Most Tm4 mistargeted at lo1 and proximal lop. (j) Tm4 targeting in *NetA* mutants is normal. (k) Tm4 targeting in *NetB* mutants is disrupted. (l) Tm4 targeting in indiffusible NetB (*NetB™*) mutants is abnormal. Most Tm4 stall at lo2-lo3. (p-s) 24% APF brains. (p-p’) NetB (green in p/gray in p’) is enriched in lobula neuropil. (q) All Tm9 (purple) arrive at lobula. Tm9 terminals at proximal lobula show the mature cylinder shape and are not colocalized with NetB (green). Tm9 terminals at distal lobula (outlined) show the bushy growth cone morphology and are colocalized with NetB. (q’) Magnification of the outlined area in q. (r) Not all Tm3 (purple) arrive at lobula. All arrived Tm3 show the bushy growth cone morphology. Therefore, Tm3 target to lobula later than Tm9. (s) Indiffusible NetB (labeled by Myc tag) are found in all neuropils. (t-w) 48% APF brains. (t) NetB (green in t/gray in t’) is enriched in M3, lo2 (yellow arrow), lo4-lo6. Lo4 is marked by the white arrow. (u) All Tm9 (purple) show cylinder-like terminals at lo1(purple arrow). Note: this driver also labels non-Tm9 neurons. (v) All Tm3 (purple) arrive at lobula and show two arbors at lo2 and lo4, which colocalize with high NetB signal. (w) Indiffusible NetB do not enrich at specific layers. Scale bar: 20μm.

Moreover, we checked the requirement of Trim9, a ring finger domain-containing E3 ubiquitin ligase functioning downstream of the Fra receptor^40–45^, in the targeting of Tm3 and Tm4. Like Fra, we found Trim9 was widely expressed in the optic lobe (Supplementary Fig. 2i). Knocking down Trim9 specifically in Tm3 or Tm4 resulted in similar axon targeting defects as knocking down Fra (Supplementary Fig. 2l, n). The knockdown efficiency of UAS-trim9 RNAi was confirmed by anti-Trim9 staining (Supplementary Fig. 2j). Thus, although Fra is expressed in diverse Tm neurons, knocking down fra in individual Tm types did not affect axons of Tm1 or Tm9 to target at lo1 but affected targeting of Tm3 and Tm4 axons to deep lobula layers (lo4). Our results are also consistent with that Trim9 mediates the downstream signaling of Fra in Tm3 and Tm4 axon targeting. Since Tm axons project to lobula in bundles, the lack of phenotype when Fra is knocked down in Tm1 orTm9 does not preclude the possibility that Fra is required collectively in Tm bundles to target the lobula, and we will address this in later sections.

Next, we checked if turning off unc-5 expression was necessary for Tm targeting to lobula. By overexpressing unc-5 with cell-type-specific drivers, we found Tm1, Tm9, Tm3, and Tm4 were mistargeted to superficial lobula layers or lobula plate (Fig. 2c, i and Supplementary Fig. 2c, f). Therefore, turning off the repulsive unc-5 is critical for properly targeting of Tm neurons to the lobula.

### Diffusible NetB enriched at deep lobula layers is required for Tm3 and Tm4 layer-specific targeting

Since Fra is required cell-autonomously for the axon targeting of Tm3 and Tm4 to lo4, we next checked whether Fra’s ligands, NetA and NetB, are required for their layer-specific axon targeting. In *netA* mutants, Tm3 and Tm4 targeting were normal (Fig. 2d, i). In contrast, in *netB* mutants, 55.4% of Tm3 and 60% Tm4 axons were stalled at lo2-lo3 (Fig. 2e, k), indicating that netB is necessary for Tm3 and Tm4 targeting layer specificity, while netA is not. Considering that Fra is required for Tm3 and Tm4 axons to target deep lobula layers but not for Tm1 or Tm9 to lo1, we wondered whether netB distribution is different in deep lobula layers vs. lo1. Therefore, we traced Tm3 and Tm9 axon targeting and NetB distribution pattern through development.

At 24% APF, NetB is distributed almost throughout the whole lobula neuropil, with higher concentration at superficial layers than deep layers (Fig. 2p, p’). At this stage, all born Tm9 axons have arrived at lobula (Fig. 2q), and most of them already have the cylinder-like axon terminals as in the adult stage. In contrast, axons at the youngest (distal) columns exhibit the bushy growth cone structure (Fig. 2q’), indicating that active axon targeting occurs in these young columns. In contrast to Tm9, a larger fraction of Tm3 axons have not arrived at lobula, and those who have arrived all still show the bushy growth cone structure (Fig. 2r). Therefore, Tm9 and Tm3 target lobula at different paces, with Tm9 targeting earlier than Tm3 in each lobula column. For old Tm9, their cylinder-like axon terminals do not colocalize with NetB, while for young Tm9, their growth cones colocalize with NetB (Fig. 2q). Similarly, all Tm3 growth cones are localized in high NetB layers (Fig. 2r). Therefore, both Tm3 and Tm9 growth cones colocalize with enriched NetB during axon targeting. At 48% APF, NetB was distributed into two regions in the lobula, one thin layer at lo2 and one big domain at lo4-lo6 (Fig. 2t, t’). All Tm9 axons exhibit the cylinder-like axon terminals and terminate at lo1 (Fig. 2u), which is devoid of NetB. For Tm3, all axons have arrived at lobula and they have terminal arborizations at two lobula layers, lo2 and lo4, both of which are enriched for NetB (Fig. 2v). Therefore, Tm3 layer-specific targeting coincides with NetB distribution, and their axon terminals colocalize with high NetB. Taken together, we found that Tm axons target lobula neuropil at different pupal stages, and early-born Tm9 axons arrive at lobula earlier than late-born Tm3 axons. Our observations suggest that during initial axon targeting, growth cones of Tm bundles extend towards high NetB concentration in lobula and Fra may be required collectively in Tm bundles but not in individual Tm types. After arriving, lobula layers are gradually refined, as different Tm growth cones reside at specific layers and become the mature axon terminal. Tm9 axons terminate at lo1 which is devoid of NetB, consistent with our observation that their layer-specific targeting is not affected by Netrin signaling. In contrast, Tm3 layer-specific targeting coincides with NetB enrichment, explaining why it is regulated by netrin signaling.

Since NetB is a secreted protein, we further checked whether its diffusibility is required for its distribution and subsequent Tm3 and Tm4 targeting. We checked if NetB distribution was affected in *netB™* flies in which NetB cannot diffuse because it was attached to a membrane-bound domain^46^. At 24% APF, NetB™ is localized in lobula and the proximal lobula plate (Fig. 2s), compared with the lobula-only restricted NetB distribution in wild type. At 48% APF, NetB™ is broadly localized to the whole lobula and part of lop (Fig. 2w). Moreover, we found about 70% of Tm3 and Tm4 were mistargeted either at lo2-3 or lop in adult *netB™* flies (Fig. 2f, l). In addition, some Tm3 axons do not join the inner chiasm but instead seemingly cut through the lobula, and we will discuss this further in later sections. In summary, NetB is enriched in deep lobula layers during Tm layer-specific targeting; and is specifically required for Tm3 and Tm4 targeting to lo4.

### Fra is required non-cell-autonomously for Tm3 axons to leave the medulla neuropil

When examining Fra’s function in Tm3 targeting, we also checked Tm3 targeting using MARCM with a well-studied loss-of-function fra allele, *fra^3^* (ref3). In addition to mistargeting to lo2 and lop, 15.6% of Tm3 stalled at medulla layer 10 (M10) (Fig. 3b, g, h). Consistently, we found in *netrin* double mutant *netAB^Δ^*, 76.7% Tm3 axons did not leave the medulla neuropil, and the integrity of lobula was disrupted (Fig. 3d, g, h). To better view the morphology defects of Tm3, we labeled individual Tm3 using MultiColor FlpOut (MCFO)^14^. We found most Tm3 axons did not innervate lobula in *netAB^Δ^* mutants. Instead, they were stalled at M10 and turned back to the medulla, forming the hook-like structure similar to Mi1 (Fig. 3f, g). In contrast, axons of Tm1 and Tm9 still leave the medulla neuropil in *netAB^Δ^* mutants (data not shown), indicating that Tm3 is specifically regulated by netrin signaling in its axonal leaving of the medulla neuropil.

**Figure 3.**
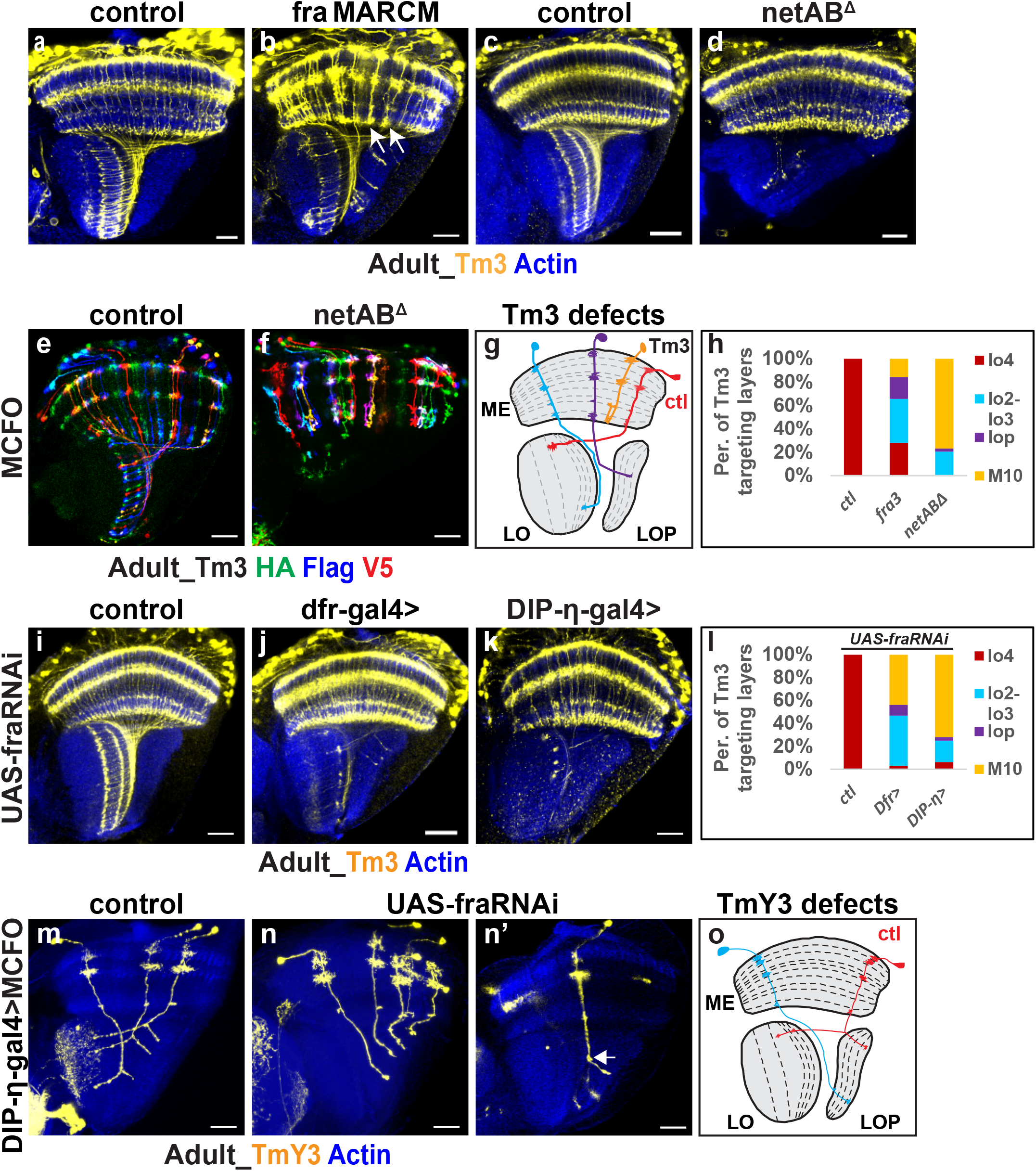
Fra acts non-cell-autonomously for Tm3 to leave the medulla neuropil. MARCM is used to label Tm3 within recombined control clones (a) or *fra* mutant clones (b). Note: Recombination efficiency is high. Tm3 stalled at M10 are marked by white arrow in (b). (c-d) Tm3 are labeled by *Tm3-lexA>lexop-tdTomato* (yellow). (c) Wild-type Tm3. (d) Tm3 targeting in *netAB* double mutants. Most Tm3 does not leave medulla neuropil, and lobula integrity is disrupted in *netAB* mutants. (e-f) MCFO is used to label individual Tm3. (e) Wild-type Tm3. (f) Tm3 in *netAB* mutants. Most Tm3 stop at M10 and their axons turned back to medulla, forming the hook structure. (g) Schematic of wild-type Tm3 (red) and abnormal Tm3 (all other colors) shown in (b) (d) (f). (h) Quantification of (b) and (d). (i-k) Tm3 are labeled by *Tm3-lexA, lexop-tdTomato*. (i) Wild-type Tm3. (j) Knocking down Fra by *Dfr-gal4* results 44% of Tm3 stalled at M10. (k) Knocking down Fra by *DIP-η-gal4* results 72% of Tm3 stalled at M10. (l) Quantification of (i-k). (m-n’) MCFO is used to label individual TmY3. *DIP-η-gal4* are expressed in various neurons, including TmY3. TmY3 is identified based on morphology. (m) Wild-type TmY3. Each TmY3 shows one lobula branch and one lobula plate branch. (n) Knocking down fra results in a tiny lobula branch, while the lobula plate branch is normal. (n’) Tiny lobula branch marked by white arrow (o) Schematic of wild-type TmY3 (red) and the TmY3 defect in (n-n’) (blue). Scale bar: 20μm.

Tm3 axons leave medulla neuropil normally when Fra is only knocked down in Tm3 specifically. In contrast, most of them do not leave medulla in *netrin* mutants, and a small percentage do not leave medulla in *fra* MARCM mutants. We hypothesized that Tm3 axons surrounded by big *fra* mutant clones were stalled at M10, meaning that Fra is required non-cell-autonomously in surrounding cells to help Tm3 axons leave the medulla. To test this hypothesis, we checked Tm3 targeting by knocking down Fra with Gal4 drivers expressed in multiple cell types. We found 44% of Tm3 neurons were stalled at M10 when Fra was knocked down using *dfr*-gal4 (Fig. 3j, l), and 72% of Tm3 were stalled at M10 when using *DIP-eta*-gal4 (Fig. 3k, l), suggesting that neurons that helped Tm3 leaving medulla were within these gal4-expressing cells. In addition, knocking down the downstream Trim9 with *dfr-*gal4 shows similar defects as knocking down Fra (Supplementary Fig. 3b).

A previous study showed that dfr+ neurons included Mi10b, Tm3a, Tm3b, Dm8a, Dm8b, Tm9, TmY3, Tm27, and Tm27Y^37^. DIP-eta-gal4 was expressed in Mi9, Mi10, Tm2, Tm3, Tm5c, TmY3, TmY13^47^. Combining these data, only Mi10, Tm3, and TmY3 are labeled by both gal4. Since Mi10 axons do not leave the medulla, TmY3 is likely essential for Tm3 leaving the medulla. If TmY3 guides Tm3 to leave the medulla, the axon guidance of TmY3 should also be regulated by Fra. Therefore, we checked the targeting of TmY3 without Fra. Since TmY3-specific driver is not available currently, we labeled individual TmY3 by crossing DIP-eta-gal4 with MCFO. Based on the arborization patterns in the medulla, we can distinguish each neural type. We found Tm2 and TmY3 were the most frequently labeled neurons (Fig. 3m, Supplementary Fig. 3c). In wild type, each TmY3 axon bifurcates into two branches, with one branch turning toward the lobula and the other project to lobula plate (Fig. 3m, o). Interestingly, after knocking down Fra, the lobula branch of TmY3 becomes tiny and pointing to lop or completely missing (Fig. 3n’, o), while the lobula plate branch and medulla arborization remain normal (Fig. 3n, n’, o), indicating that Fra is specifically required for the lobula branch of TmY3 to target the lobula neuropil.

Moreover, we found Tm2 left medulla and targeted normally when Fra is knocked down with DIP-eta-gal4 (Supplementary Fig. 3d). In summary, our results suggest that Tm3 targeting can be divided into distinct steps, first leaving the medulla neuropil and then joining the IOC. Fra is required non-cell autonomously for Tm3 to leave the medulla neuropil. Moreover, Fra is necessary for TmY3 to extend a lobula branch which likely helps Tm3 leave the medulla neuropil.

### Fra expressed by medulla neurons enrich NetB in the lobula

We showed that NetB diffusibility is essential for Tm3 targeting to lo4, and at 48% APF, NetB was enriched at two domains, lo2 and lo4-lo6. Next we asked how the diffusible NetB was enriched in these two separate domains. Previous studies found that Fra receptors expressed on axon terminals localize and enrich Netrin in embryonic CNS^48^ and medulla layer M3^25^. We examined whether Fra also regulates NetB distribution in the lobula. First, we checked the distribution pattern of Fra and NetB in neuropils through development. At 24% APF, we found that Fra and NetB colocalize throughout the lobula neuropil (Fig. 4a). At 48% APF, netB was enriched at restricted domains, still colocalizing with Fra (Fig. 4b). Next, we checked if NetB distribution was affected without Fra. We knocked down Fra by crossing *fraRNAi* with *ap-gal4* (*ap>fraRNAi* hereafter), a gal4 covering about half of the optic lobe neurons, including the majority of Tm and TmY neurons^13,31^. At 24% APF, NetB level in the lobula is reduced and NetB at the cleft between neuropils is increased in *ap>fraRNAi* comparing with control (Fig. 4c-d’, h). NetB signal in cell bodies is normal in *ap>fraRNAi*, indicating that NetB synthesis is not affected (Fig. 4c-d’). At 48% APF, in contrast to NetB enriching at lo2 and lo4-6 in wild type, lo2 NetB was almost missing, and lo4-lo6 was significantly reduced in *ap>fraRNAi* (Fig. 4f, i). Moreover, NetB enrichment at M3 is continuous though the level is reduced. Interestingly, we also observed disrupted inner chiasm (shown by Tm9 axons not joining the inner chiasm) and irregular lobula structure in *ap>fraRNAi* (Supplementary Fig. 4b) and will discuss this later. Because ap-gal4 also labels a subset of neurons generated from the IPC, we knocked down Fra using SoxN-gal4, which is expressed only in the main medulla region neuroblasts and subsequently inherited in medulla neurons. Like ap>fraRNAi, NetB level in lobula is reduced in *SoxN>fraRNAi* comparing with control (Fig. 4g, i). Moreover, NetB level at M3 is greatly reduced, consistent with targeting area Fra helps enrich NetB at M3 found in a previous study^25^. To sum up, these data indicated that Fra co-localized with NetB in neuropils, and Fra expressed by medulla neurons (Tm/TmY neurons) contributes to NetB enrichment in the lobula.

**Figure 4.**
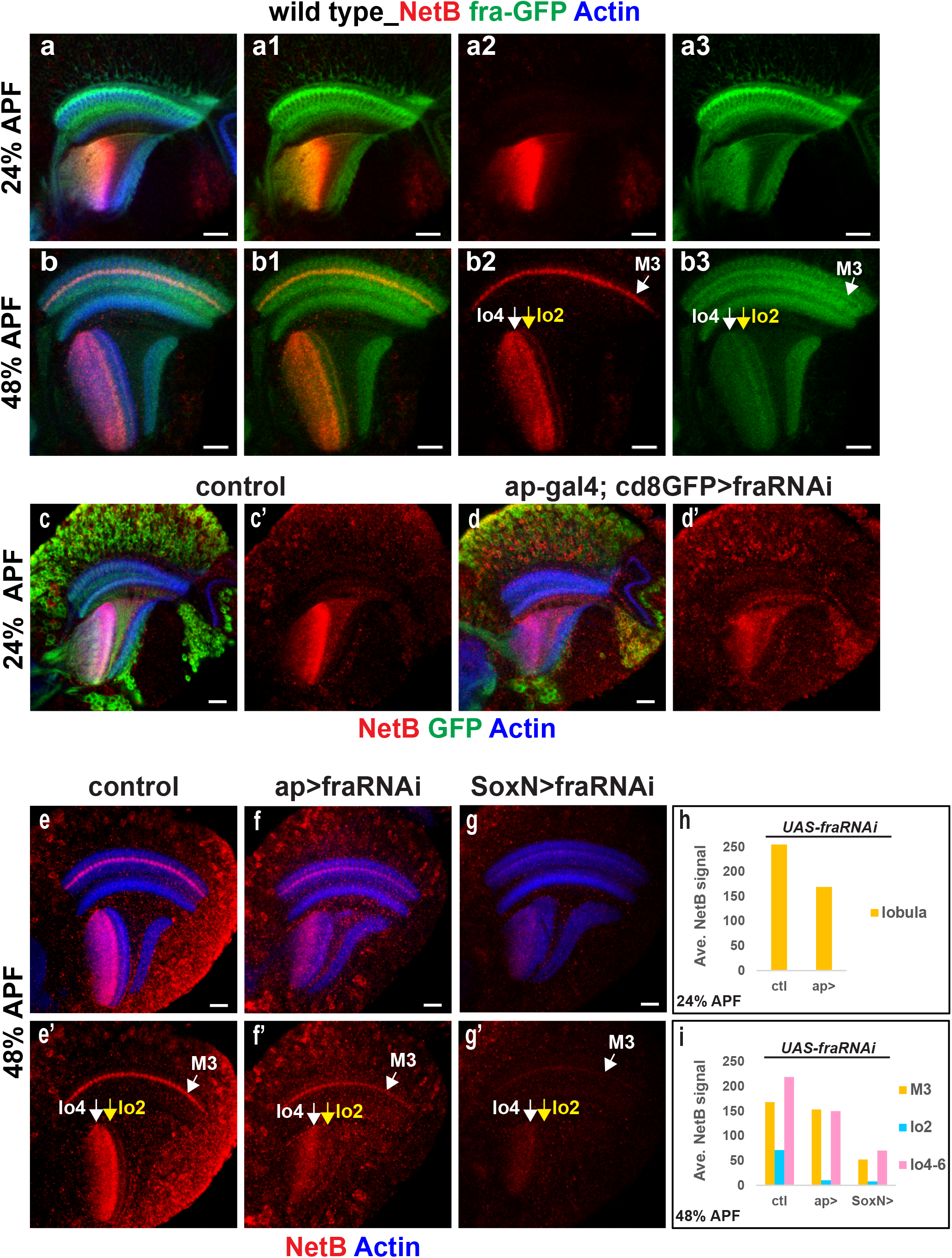
Fra from medulla neurons regulates NetB distribution. (a) Fra (green) and NetB (red) are costained at 24% APF. Fra and NetB are colocalized at lobula. Actin is used to label all neuropils. (b) Fra (green) and NetB (red) are costained at 48% APF. All NetB enriched areas (M3, lo2, lo4-lo4) are co-localized with Fra. (c-d) 24% APF brains are shown. (c) Control. *Ap>cd8GFP* (green) labels half of the neurons in optic lobe. (d) Knocking down Fra by *ap-gal4*. NetB signal in medulla cortex is normal, indicating NetB synthesis is not affected. The NetB level at lobula dropped comparing with control. (e-g) 48% APF brains are shown. (e) Control. NetB enriched layers are shown. (f) Knocking down Fra by *ap-gal4*. M3 NetB level drops but still continuous. Lo2 NetB is almost missing. Lo4-lo6 NetB is greatly reduced. (g) Knocking down Fra by *SoxN-gal4*, which are expressed in medulla neuroblast and inherited by progenies. M3 NetB is greatly reduced. Lo2 NetB is almost gone. Lo4-lo6 NetB is greatly reduced. (h) Quantification of (c-d). (i) Quantification of (e-g). Scale bar: 20μm.

### Unc-5 regulates NetB distribution

Since not all Fra-enriched lobula layers in the optic lobe showed NetB distribution, we hypothesized other molecules should also contribute to NetB distribution. The other known NetB receptor, Unc-5^35^, is a good candidate because we found that the distribution patterns of Unc-5 and NetB in optic lobe neuropils were always mutually exclusive with each other in the early to mid-pupal stages. At 24% APF, Unc-5 is enriched in the distal medulla layers, the posterior young medulla columns and all lobula plate layers (Fig. 5a2). In contrast, NetB is only enriched in lobula neuropil where no Unc-5 resides (Fig. 5a1). At 36% APF, Unc-5 and NetB distribution are almost the same as at 24% APF except that Unc-5 does not enrich at the posterior medulla columns since all columns have formed at this stage (Supplementary Fig. 5a). At 48% APF, Unc-5 is highly enriched in the distal medulla layers, proximal medulla layers, lo1, and all lop layers (Fig. 5b2). In contrast, NetB is enriched in M3, lo2, lo4-lo6 (Fig. 5b1). At 60% APF, Unc-5 at lo1 is reduced, and NetB signals at lobula are all gone (Supplementary Fig. 5b). Therefore, during the whole axon targeting period of optic lobe neurons, Unc-5 does not colocalize with NetB, and NetB is localized to layers where there is Fra but no Unc-5.

**Figure 5.**
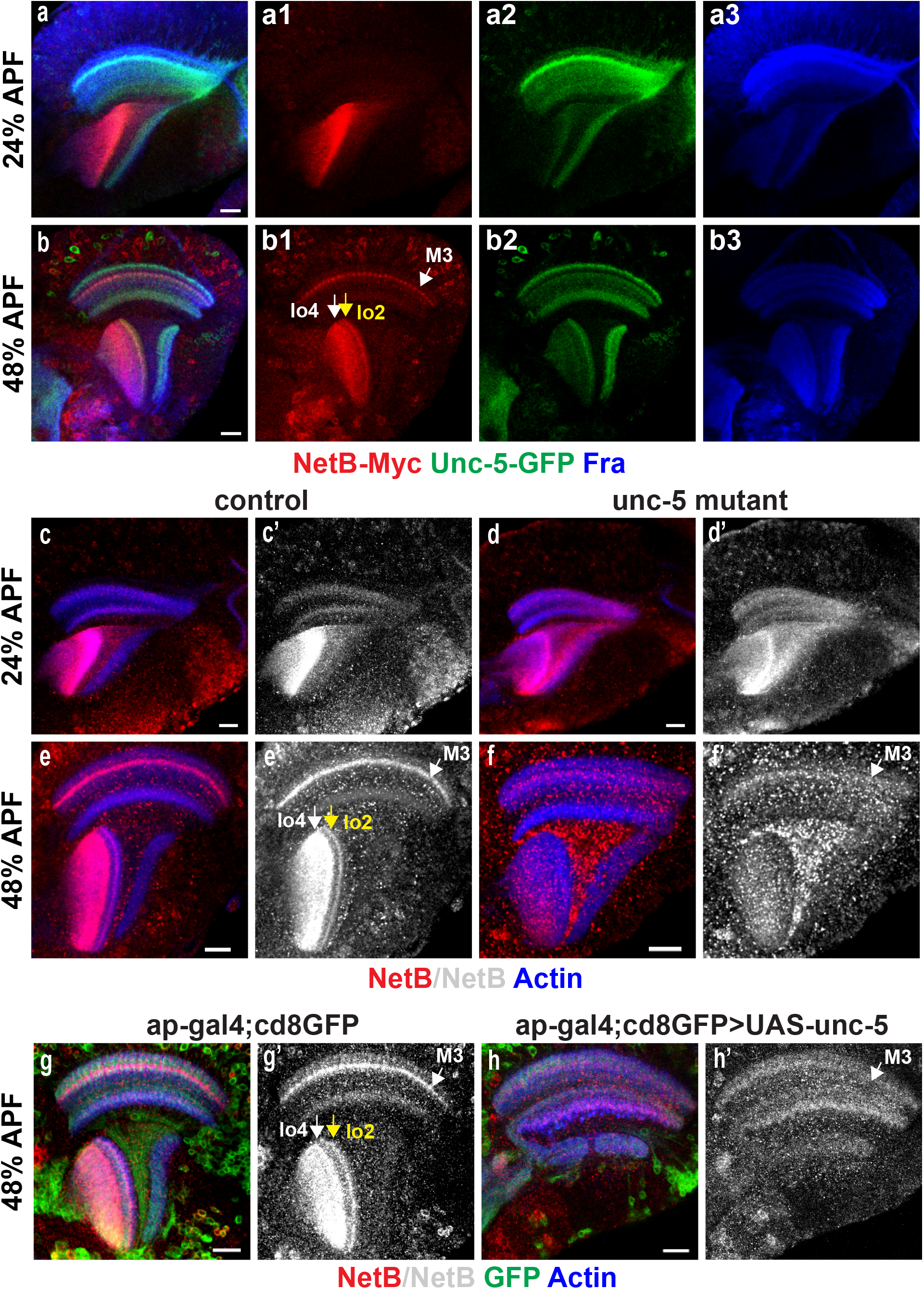
Unc-5 regulates NetB distribution. (a) Unc-5 (green), Fra(blue) and NetB (red) are costained at 24% APF. Unc-5 and NetB are not colocalized. (b) 48% APF brains. Unc-5 and NetB are not colocalized. (c-d) 24% APF brains. (c-c’) NetB (red and gray) distribution in wild-type. (d-d’) NetB distribution in *unc-5* mutants. NetB also shows at lobula plate. (e-f) 48% APF brains. (e-e’) NetB distribution in wild-type. (f-f’) NetB distribution in *unc-5* mutants. M3 NetB (white arrow) is almost gone; lobula NetB level is greatly reduced; intense NetB signal shows at the cleft between neuropils. (g-h) 48% APF brains. (g-g’) NetB distribution in wild-type. (h) Overexpressing Unc-5 by *ap-gal4*. Lobula complex is severely damaged. No NetB is enriched at lobula. M3 NetB (white arrow) is greatly reduced. Actin is used to label all neuropils. Scale bar: 20μm.

Next, we examined whether Unc-5 contributed to NetB distribution. In *unc-5* null mutants^49^, we found that the NetB signal in the cell bodies at the medulla cortex was the same as the wild type (Fig. 5d,d’), demonstrating that NetB synthesis was normal in *unc-5* mutants. However, at 24% APF, NetB is spreading to both lobula and lobula plate in *unc-5* mutants (Fig. 5d, d’). At 48% APF, NetB is no longer enriched in specific layers in lobula; instead, high-intensity dotted signals of NetB are observed in the cleft among neuropils (Fig. 5f, f’). Moreover, the NetB signal at M3 layer was also smearing in *unc-5* mutants, indicating that Unc-5 affected NetB distribution throughout the optic lobe. We also checked Tm3 targeting without Unc-5 and found that Tm3 was mistargeted to lo2 or lop in unc-5 mutants (Supplementary Fig. 5d).

Consistently, Tm3 was also mistargeted to lo2 or lop when unc-5 is knocked down in all neurons with elavGal4>unc5RNAi (Supplementary Fig. 5f). In addition, the inner chiasm and the integrity of lobula were disrupted in *unc5* mutants (Supplementary Fig. 5h), as in *ap>fraRNAi*. Therefore, Unc-5 regulates NetB distribution, and loss of Unc-5 causes NetB to spread abnormally.

Next, we checked whether ectopically expressing Unc-5 would affect NetB enrichment. We overexpressed unc-5 by crossing *UAS-HA-unc5* with *ap-gal4*. Our data showed that with overexpression of unc-5, the NetB level at M3 was significantly reduced (Fig. 5h, h’). In addition, the structure of the lobula complex was totally damaged; lobula and lobula plate can be barely distinguished, and no enrichment of NetB was observed in any part of the lobula complex (Fig. 5h, h’). Taken together, Unc-5 is necessary for NetB distribution, and ectopic expression of Unc-5 is sufficient to affect NetB enrichment.

### Netrin signaling regulates the formation of the inner chiasm

Previously, we mentioned that in *netB™*, *ap>fraRNAi* and *unc-5* mutants, the IOC was disrupted shown by Tm3 or Tm9, indicating that Netrin signaling also contributes to IOC formation. In the fly optic lobe, to maintain the retinotopic information, the anterior-most medulla columns always connect with the proximal-most lobula columns and the posterior-most medulla columns with the distal-most lobula columns (Fig. 1a). The X-shaped IOC is formed since posterior axons need to cross all the anterior axons before reaching the corresponding lobula columns. The main components of IOC include Tm, TmY, and T2/T3 neurons^21^. Tm axons first project through the medulla column, join the IOC, then enter the corresponding lobula column, and finally terminate at specific lobula layers. As previously shown, Tm3 marked IOC was not affected in either *netA* mutants or *netB* mutants (Fig. 2d, e), indicating that NetA and NetB should act redundantly for IOC formation. Therefore, we checked IOC formation in *netAB* double mutants. We found that IOC labeled by Tm1, Tm9, or Tm2 axons was all disrupted in *netAB* mutants (Fig.6e, f, g), despite their axons still terminated in correct layers. Tm4 axons left medulla neuropil, but they did not form normal IOC and terminated at wrong lobula layers (Fig.6h). Taken together, these data suggested that NetA and NetB act redundantly in IOC formation, and diffusibility of Netrins is also required since *netB™* brains also have IOC defects (Fig. 2f).

**Figure 6.**
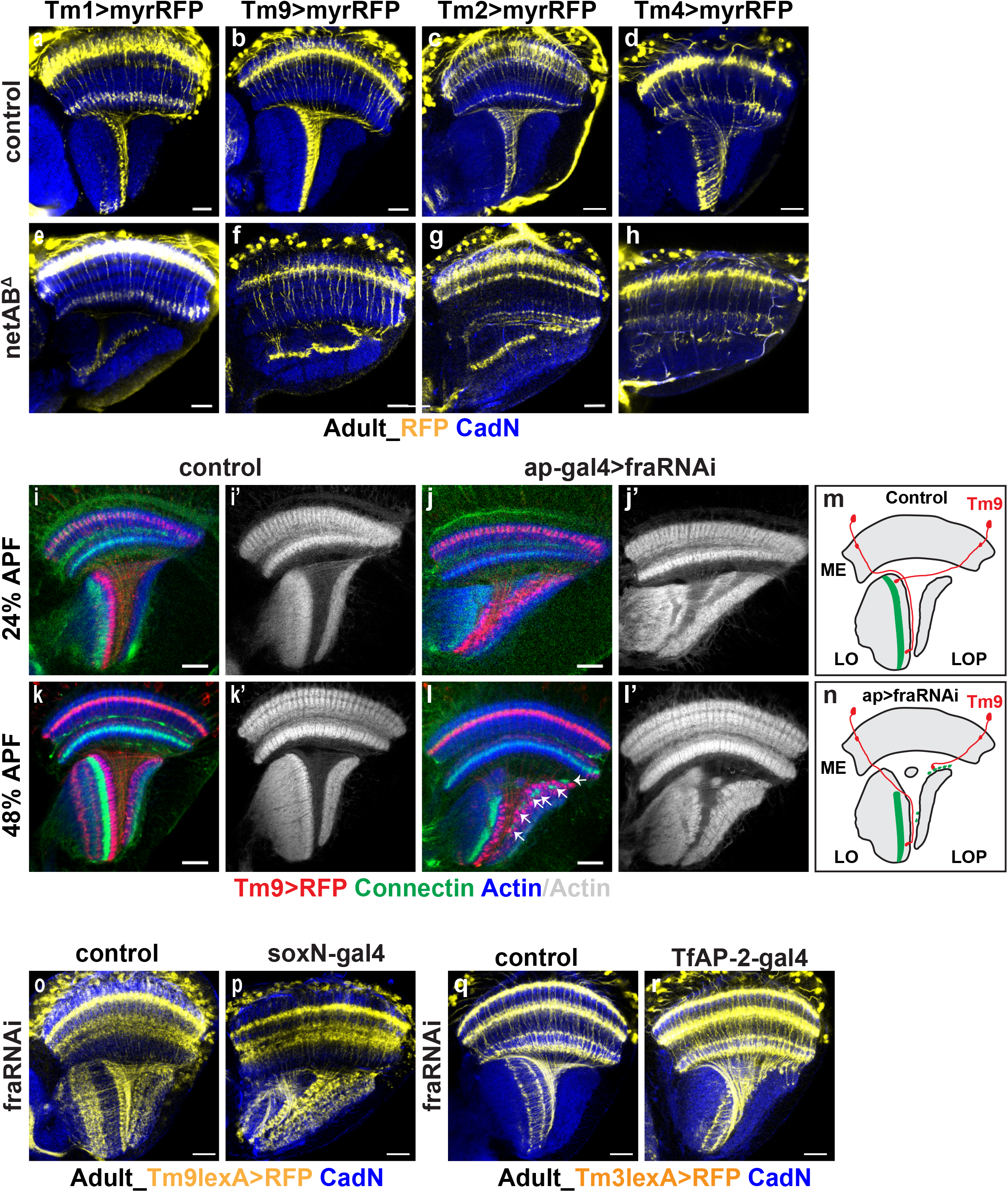
Netrin signaling regulates IOC formation. (a-e) Tm1, Tm9, Tm2, Tm4 in wild-type. (e-h) Tm1, Tm9, Tm2 and Tm4 in *netAB* mutants. IOC is disrupted. Tm1, Tm9 and Tm2 layer-specific targeting are not affected. Tm4 are mistargeted. CadN (blue) labels all neuropils. (i-j’) 24% APF brains growing at 29°C. (i) Tm9 (red) in wild-type. Axons terminate at lo1. Connectin (green) labels middle layers of lobula. (i’) Neuropils (gray) are intact. (j) Tm9 (red) in *ap>fraRNAi*. Axons are mislocated. Connectin (green) band is scattered. (j’) Lobula (gray) organization is damaged. (k-l’) 48% APF brains growing at 29°C. (k) Tm9 (red) in wild-type. Axons terminate at lo1. Connectin (green) labels lo3 and distal half of lo4. (k’) Neuropils (gray) are intact. (l) Tm9 (red) in *ap>fraRNAi*. Axons are mislocated. Note: Tm9 driver also labels other neural types. Tm9 axon terminal is cylinder-shaped. Connectin (green) band is scattered at lobula and also accompanied with mislocated Tm9 terminals. (l’) Lobula (gray) organization is damaged. (m) Schematic of Tm9 in wild-type. Anterior Tm9 (blue) terminate at proximal lobula, and posterior (Red) terminate at distal lobula, forming X-shapted IOC. Connectin band is shown in green. (n) Schematic of Tm9 in *ap>fraRNAi*. Tm9 from late-born columns do not cross all the early-born columns but stop half-way. Mislocated Tm9 axon terminal still mature and are accompanied with connectin signals. (o-p) Tm9 are labeled by *Tm9lexA>lexop-tdTomato* (yellow). (o) Tm9 in wild-type. (p) Tm9 in *SoxN>fraRNAi*. IOC is disrupted. Tm9 axon terminal mature normally. Therefore, medulla neurons are required for IOC formation. (q-r) Tm3 are labeled (yellow). (q) Tm3 in wild-type. (r) Tm3 marked IOC are damaged in *TfAP-2>fraRNAi*. Therefore, the *TfAP-2-gal4*-expressing neurons, which are born in early-temporal window, are required for IOC formation. Scale bar: 20μm.

To better understand how Netrin signaling affects Tm joining the inner chiasm, we checked Tm targeting at 24% and 48% APF when axon targeting was still undergoing. We visualized the inner chiasm using Tm9 as its axon termination layer was not affected by netrin signaling. We knocked down Fra using *UAS-fraRNAi* driven by *ap-gal4* and checked the inner chiasm formation. In wild type, at 24% and 48% APF, the lobula neuropil was intact as labeled by phalloidin; all Tm9 neurons joined the inner chiasm and terminated at the superficial lobula layer (Fig.6i, k). However, in *ap>fraRNAi* brains, some Tm9 axons do not turn to lobula but instead stop at the surface of the lobula plate; some Tm9 axons do not cross all the adjacent anterior axons but stop at halfway (Fig.6j, l). Moreover, at 48% APF, Connectin labels a continuous lobula band in wild type (Fig.6k, m). In contrast, ectopic Connectin signals appear at the surface of the lobula plate where Tm1 axon terminals are mislocated in *ap>fraRNAi*.

Consistently, lobula parts without Tm9 terminals are devoid of Connectin signals, creating the discontinuous Connectin band of lobula in *ap>fraRNAi* (Fig.6l, n). To sum up, these data suggest that Netrin signaling is required for the IOC formation. Without Netrin signaling, late-born axon bundles cannot cross all of the early-born axon bundles; instead, they stop halfway and continue the maturation locally, creating the disrupted IOC and the scattered lobula phenotype.

### Fra expressed in early-born Tm neurons is required for the formation of the inner chiasm

Since the inner chiasm comprises cells from the medulla (Tm and TmY neurons) and lobula complex (T2 and T3), we tested Fra from which part is essential for the inner chiasm formation. We knocked down the medulla Fra by crossing *UAS-fraRNAi* with *SoxN-gal4* and found that the inner chiasm was disrupted (Fig.6p). *SoxN-Gal4* is expressed only in the main medulla region neuroblasts and inherited subsequently in medulla neurons, therefore, Tm (and TmY) neurons are required for the inner chiasm formation. To examine whether there is a specific subset of Tm neurons crucial for IOC formation, we first used Gal4 drivers initiated in different temporal windows to knock down Fra. Medulla neuroblasts sequentially generate different neural types controlled by a temporal transcription factor (TTF) cascade: Hth -> Opa -> Ey+Erm -> Ey+Opa -> Slp -> D -> B-H1&2->Tll->Gcm^17^. We found knocking down fra using *ey-gal4*, *slp-gal4*, or *D-gal4* does not affect the inner chiasm formation (Supplementary Fig.6b, c, d), indicating that Fra expressed in Tm neurons born in after the Ey window are not necessary for the formation of the inner chiasm. An earlier TTF Opa is required for the generation of TfAP-2 expressing neurons, which include Tm1, Tm2 and Tm4^18,31^. We found a TfAP-2 driver (GMR24B02-GAL4) that shows expression at 3^rd^ instar larval stage in the Flylight Gal4 collection. When Fra was knocked down by TfAP-2-gal4, the IOC formation was disrupted (Fig.6r). As previously shown, knocking down Fra specifically in Tm1 or Tm4 did not cause a defect in IOC formation (Supplementary Fig. 2b, Fig.2h), indicating that loss of Fra from individual Tm type does not affect the IOC formation. To sum up, we found that Fra expressed in a group of early-born Tm neurons are collectively necessary for the IOC formation, and these neurons likely include TfAP expressing Tm1, Tm2, and Tm4 neurons that are generated in the early (Opa) temporal window.

### Notch signaling regulates NetB expression

Neuron identities are determined by neural fate specification programs, which then induce the expression of downstream effectors such as guidance cues and cell surface molecules. To connect NetB expression with the upstream regulator, we analyzed scRNA-seq data of early pupal optic lobe neurons^31^ and found that the top transcription factor whose expression correlates with NetB is Apterous (Ap). Ap is expressed in all Notch-on medulla neurons, and Ap expression is lost when Notch signaling is lost in *Su(H)* mutant clones^15^. We checked if NetB was expressed by Notch-on neurons and found that indeed all of the NetB expressing cells in the medulla are also labeled by Ap-lacZ (Fig. 7a). Next, we examined whether Ap regulated NetB expression. However, NetB expression was not affected in *ap* mutants (Supplementary Fig. 7a). Then, we checked if NetB expression was regulated by Notch signaling and found that NetB signal was lost in *Su(H)* mutant MARCM clones (Fig. 7d). Consistently, knocking down Su(H) by elav-gal4> Su(H) RNAi abolished NetB expression in the medulla cortex (Supplementary Fig. 7c). Taken together, these data suggest that Notch signaling is required to activate NetB expression in medulla neurons.

**Figure 7.**
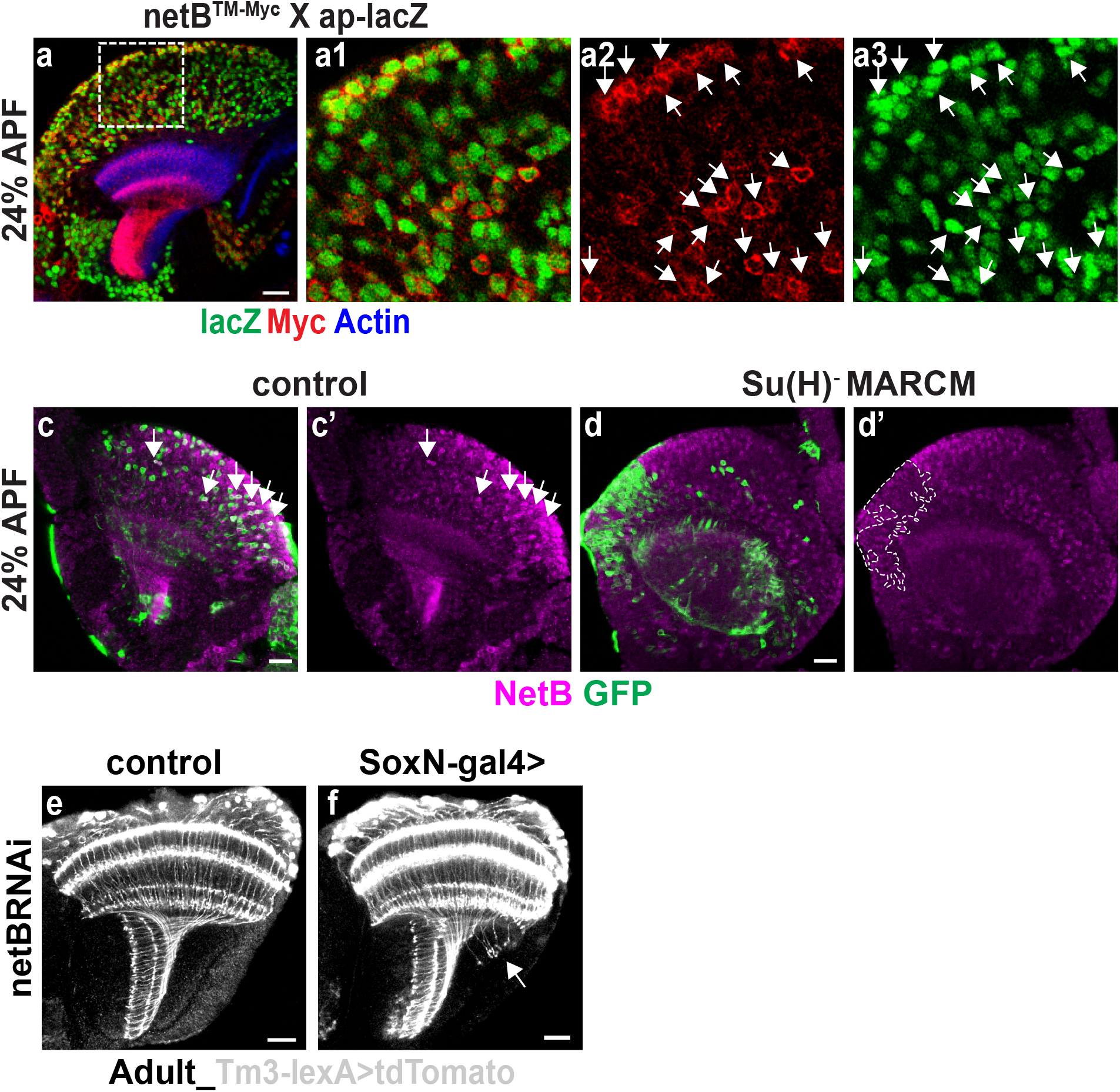
Notch regulated NetB expression at medulla. (a-a3) NetB (red) is costained with ap-lacZ (green), which is activated in Notch-on neurons at medulla. All NetB-expressing medulla neurons are Notch-on. Actin (blue) labels neuropils. (c-d’) MARCM is used to create recombinated control clones (c) or mutated *Su(H)* clones (d). (c) In control, some clones colocalize with NetB signals (white arrow). (d) Su(H) clones do not show any NetB signal. (d’) Big *Su(H)* clones are outlined. (e-f) Tm3 are labeled (gray). (e) Tm3 in wild-type. (f) Tm3 in *SoxN>netBRNAi*. *SoxN-gal4* are expressed in medulla neuroblasts and inherited by progenies. Knocking down NetB in medulla only affect the Tm3 from late-born columns, suggesting that medulla originated NetB contributes to IOC formation. Scale bar: 20μm.

### Medulla-secreted NetB contributes to the joining of late-born axon bundles to IOC

Tm neurons from anterior medulla columns arrive at the lobula earlier and contribute to the NetB pool^25^, which might influence Tm targeting from posterior columns that arrive later. To test whether NetB derived from medulla neurons affects Tm targeting in later-born posterior columns, we knocked down NetB using SoxN-gal4>netBRNAi and observed disrupted IOC formation from Tm neurons in posterior columns (Fig. 7f). Taken together, the Notch-dependent binary fate choice regulates the ligand of the Netrin guidance pathway and contributes to the correct assembly of the neuropils.

## Discussion

### Netrin signaling has multiple roles at different steps of Tm axon targeting

The fly optic lobe consists of repeated columnar and layered units. Medulla columns connect with lobula columns in a stereotypical manner, forming a retinotopic map to maintain the visual information. Anterior medulla columns are born early and connect with the proximal lobula columns. Posterior medulla columns are born late and connect with the distal lobula columns. After leaving the medulla column, posterior bundles need to cross all of the anterior bundles before arriving at the lobula, forming the X-shaped IOC. One of the main components of IOC are axons of Tm neurons. Tm axons extend through a medulla column, join the IOC to target lobula, enter the corresponding lobula column, and terminate at a specific layer. After Tm bundles leave the medulla, they have three directions to choose from: turning anterior to target lobula, turning posterior to target lobula plate, and going forward along the cleft between lobula and lobula plate. Netrins are enriched only in lobula but not in lobula plate. The attractive receptor Fra is expressed in Tm neurons while the repulsive receptor Unc-5 is not, making Netrin signaling a good candidate for regulating Tm targeting. We show that Netrin signaling is required for Tm bundles to choose between lobula and lobula plate, cross the anterior bundles to target lobula thus forming the IOC, and also for the layer specificity of a subset of Tm neurons that terminate at deep lobula layers. NetA and NetB act redundantly in the IOC formation, and Netrin diffusion is required as well, while the layer-specific targeting of Tm3/Tm4 primarily relies on NetB. Netrin pool in the lobula is contributed by different sources, including lobula neurons, T2 neurons and medulla Tm neurons. Our results suggest that Tm bundles from early born (anterior) medulla columns are guided by lobula netrin and after they arrive, they provide more Netrins to the lobula pool to facilitate Tm axons from late-born (posterior) columns to target lobula.

Netrin signaling is not required for Tm axons to extend forward along the cleft until they reach their retinotopic targeting position, since the retinotopic map is preserved in netrin mutants. Therefore, other mechanisms exist to control this aspect of Tm targeting.

### Early-born Tm neurons pioneer the IOC and help organize superficial lobula layers

There are about thirty types of Tm neurons, which are born sequentially in different temporal windows^15^. Do Tm axons target collectively or separately? Knocking down fra in Tm1 or Tm9 specifically does not affect their joining the IOC, but knocking down fra in broader neuron types can affect them to join IOC, indicating that Tm neurons migrate collectively towards lobula. Tm bundles include various Tm types. However, not all of them equally contribute to the IOC formation. We found that knocking down Fra in Tm neurons born in Ey stage onward does not affect IOC formation, while loss of Fra in early-born TfAP-2-expressing Tm neurons cause defects in IOC formation. Although knocking down fra in all Tm neurons results in more severe IOC defects, our results still suggest that the TfAP-2-expressing early-born Tm neurons are likely to be the pioneering axons for the initial establishment of IOC.

Without Netrin signaling, Tm axons that fail to cross anterior Tm bundles terminate at the surface of lobula plate or the IOC region but not at the distal lobula. However, mislocated Tm axons still form normal terminal arborizations. Furthermore, the following layer maturation continues around the mislocated Tm terminals, leading to discontinuous lobula superficial layers as shown by connectin labeling, creating the appearance that adjacent anterior Tm axons seem to be cutting through the lobula neuropil. Therefore, our results suggest that loss of Netrin signaling affects the proper location of distal lobula columns by disrupting IOC, but the maturation of distal lobula layers is not affected by loss of netrin signaling or proximal lobula layers.

### Tm3 has a special requirement for Netrin signaling to leave the medulla

In *netrins* double mutants, most Tm3 axons do not leave the medulla neuropil while axons of other Tms, including Tm1, Tm2, Tm9, and Tm4 neurons do leave. Therefore, Netrin signaling is specifically required for the leaving of Tm3 axons from the medulla before joining the IOC. In this process, NetA and NetB act redundantly, and diffusion of ligands is not required. Moreover, Fra is not needed cell-autonomously for the leaving of Tm3 axons from medulla. Instead, Tm3 needs help from other neuron types whose targeting should also be regulated by Netrin signaling.

TmY3 is likely to be the neuron that helps Tm3 leave the medulla. Without fra, TmY3 lost its innervation of lobula, showing a tiny neurite at the branching point. How can losing the lobula branch of TmY3 affect Tm3 leaving the medulla? Since the mechanism for TmY branching has not been well understood, we guess that TmY3 branches immediately after it leaves the medulla. Therefore, without the guidance of the lobula branch of TmY3, Tm3 no longer leaves the medulla. A caveat is that Tm3 outnumbers TmY3, with a ratio of 11:4^50^. Therefore, TmY3 may not guide Tm3 to leave the medulla by physical interaction.

Tm3 neurons join IOC later than Tm9 and are not part of the pioneer bundle that organizes the IOC. Tm3 targeting is more sensitive to netrin signaling when joining IOC because knocking down fra specifically results in mistargeting to lop, which is not seen in Tm9 or Tm1. Moreover, Tm3 layer-specific targeting is regulated by diffusible NetB, and NetA does not act redundantly with NetB in this process.

### Fra and Unc-5 regulate NetB enrichment in the optic lobe

Diffusible netrins are required for IOC formation and Tm3 layer-specific targeting. Consistently, NetB shows enrichment in M3, lo2, and lo4-lo6 at the mid-pupal stage. Previous studies found Fra captures Netrins to enrich them locally^25,48^. Here we also found that Fra is colocalized with NetB in neuropil layers. Without Fra expressed in Tm neurons, NetB enrichment at lobula is greatly diminished. However, since Fra also guides the targeting of Tm neurons to lobula and Tm neurons are also part of the Netrin source, it is possible that both mechanisms contribute to the diminished NetB in the lobula.

Although fra is required for NetB enrichment, not all fra are colocalized with NetB in the optic lobe. Therefore, we checked if other molecules are involved in NetB distribution. We chose Unc-5 as a candidate since its expression was exclusive with NetB in neuropils. We found that Unc-5 affects NetB distribution. In unc-5 mutants, NetB cannot be enriched in lobula and spread to lobula plate. Consistently, IOC formation and Tm3 layer-specific targeting are disrupted in unc-5 mutants. Moreover, overexpression of Unc-5 results in disrupted NetB distribution. However, how does Unc-5 regulate NetB distribution needs future investigation. Unc-5 may affect NetB distribution directly or by affecting the guidance of specific neurons, which affect NetB distribution.

### Notch dependent regulation of Netrin expression contribute to the neuropil assembly

We showed that all NetB expressing neurons in the medulla belong to the Notch-on hemilineage, and further we found that Notch signaling is required to activate NetB expression in medulla neurons. The majority of Tm neurons belong to the Notch-on hemilineage, thus activation of NetB expression by Notch signaling ensures that lobula-targeting Tm neurons will provide more NetB to the lobula NetB pool. Tm axons from later-born posterior medulla columns may require a higher concentration of NetB to target lobula, because of the longer distance they need to migrate. Consistently, when we knocked down medulla-derived NetB using SoxN-gal4, Tm axons from posterior columns showed disrupted ioc formation. Taken together, our results suggest that the Notch-dependent binary fate choice in medulla neurons is required for the regulation of an axon guidance cue, which is required to the correct assembly of neuropils.

## Methods

### Fly stocks and genetic crosses

Fly stocks were maintained in standard medium at 25°C except for RNAi experiments, for which stocks were growing at 29 °C. Wandering 3^rd^ instar larvae were picked for larval stage immunostaining. White pupae were growing to certain age either in a Petri dish layered with hydrated filter paper when growing at 25°C or in a vial of food at 29°C. The pupal stage lasts about 100 hr when growing at 25°C and about 78hr at 29°C. As a result, 24% APF means growing white pupae for 24 hours at 25°C and about 19 hours at 29°C. The following stocks/crosses were used for expression analyses: *netA^Δ^B^myc^* (from B. J. Dickson); *fra-GFP* (BL 59835); *unc-5-GFP* (BL 64547); *UAS-myr-mRFP* (BL7118); *UAS-cd8GFP; 27b-gal4*^13^; *Tm9-gal4* (BL 48050); *Tm3-gal4*(BL48569) for adult labeling; *Tm3-gal4* (BL76324) for larva labeling; *Tm4-Gal4* (BL49922); *Tm3-lexA* (BL52459), *lexopTdTomato*; *ap-lacZ*^51^. The following stocks/crosses were used for loss of function with mutants: *netA^Δ^*, *netB^Δ^*, *netA^Δ^B™* and *netA^Δ^ B^Δ^* (from B. J. Dickson); *Unc-5^8^* (from G. J. Bashaw); *MCFO* (BL64085). The following stocks/crosses were used with UAS-RNAi: *UAS-fraRNAi* (BL 40826); *UAS-Trim9RNAi* (VDRC: 21405, 21406); *UAS-unc-5RNAi* (BL33756); *tll-gal4* (GMR31H09, BL 49694); *elav-gal4*; *UAS-Dcr2* (BL 25750); *dfr-gal4* (from M. Sato); *DIP-η-gal4* (BL90319); *ap-Gal4*^51^; *SoxN-gal4* (GMR41H10Gal4); *Tm9-lexA* (BL54982); *ey-gal4* (GMR16F10, BL48737); *slp-gal4* (GMR35H02, BL 49923); *D-gal4* (GMR12G08, BL47855).

The following stocks/crosses were used with gain-of-function analyses: *UAS-HA-Unc-5* (from G. J. Bashaw). To generate wild-type of fra mutant MARCM clones, flies of *ywhsFLP UASCD8GFP; FRT G13/Cyo; Tm3-Gal4/TM6B* were crossed with *FRTG13* or *FRTG13*, *fra^3^*(BL8813). To generate wild-type or *Su(H)* mutant MARCM clones, flies of *y,w,hsFLP,UAS-CD8∷GFP; FRT40A, tub-gal80; tub-gal4/TM6B* were crossed with *FRT40A* or *FRT40A, Su(H)^Δ47^*/*CyO* flies (gift from F. Schweisguth). To induce mutant clones, the progenies were heat-shocked at 37 °C at early larval stage and dissected at adult or 30% APF. For ap mutants, flies of Mi{FlpStop}apMI01996-FlpStop.ND/CyO (BL67675) were crossed with ap-gal4. To induce mutant clones, the progenies were heat-shocked at 37 °C at early larval stage and dissect at 30% APF^52^.

### Immunostaining and Antibodies

Immunostaining was done as described previously^15^ with a few modifications. Brains were dissected in phosphate-buffered saline (PBS), fixed for 30 min for 3^rd^ instar larvae, 40 min for pupae and adult at room temperature in 4% paraformaldehyde in PBS, washed in PBS containing 0.3% Triton X-100. Brains were then incubated in primary antibody solution at 4 °C overnight, washed three times with PBST, incubated in secondary antibody solution at room temperature for 3 hours, washed three times with PBST and three times with PBS and mounted in Slowfade.

The following primary antibodies were used: rabbit anti-NetB (from B.Altenhein, 1:100), rabbit anti-NetA (from B.Altenhein, 1:100), sheep anti-GFP (AbD Serotec: 4745-1051, 1:500), Phalloidin-iFluor 405 (Abcam: ab176752, 1:1000), rabbit anti-RFP (Abcam: ab62341, 1:1000), rabbit anti-fra (from Y.-N. Jan, 1:400), rat anti-Dfr (from M. Sato, 1:100), mouse anti-Cut (DSHB:2B10, 1:10), Chicken anti-lacZ (Abcam: ab9361, 1:800), Guinea pig anti-Toy (from C. Desplan, 1:200), rabbit anti-Sox102F (from C. Desplan, 1:100), mouse anti-Myc (DSHB: 9E10, 1:10), Guinea pig anti-Trim9 (from K. Emoto, 1:200), rabbit anti-HA (CST #3724, 1:1500), rat anti-Flag (Novus: NBP1-06712, 1:100), V5 (Bio-Rad: DyLight^®^550, 1:800), rat anti-DN-cadherin (DSHB: DN-Ex#8, 1:50), mouse anti-Connectin (DSHB:C1.427, 1:10)

Secondary antibodies were from Jackson or Invitrogen.Images are acquired using a Zeiss Confocal Microscope. Figures are assembled using Photoshop and Illustrator.

### Quantifications

From the targeting defect: at least four brains were imaged, and in each brain numbers of axon terminals from at least three different focal planes were counted. For NetB level analysis, images were taken with the same confocal parameters for control and test groups. Histogram tool from photoshop was used and median values from regions of interest were calculated.

## Acknowledgements

We thank the fly community, especially Greg J. Bashaw, Benjamin Altenhein, Yuh Nung Jan, Kazuo Emoto, Barry J. Dickson, and Claude Desplan for generous gifts of antibodies and fly stocks. We thank the Bloomington Drosophila Stock Center, the Vienna Drosophila RNAi Center, the Developmental Studies Hybridoma Bank, and TriP at Harvard Medical School (NIH/NIGMS R01-GM084947) for fly stocks and reagents. We thank Filipe Pinto-Teixeira and Isabel Holguera for discussion at the beginning of the story. This work was supported by National Institutes of Health (Grant 1 R01 EY026965-01A1).

## Author information

The authors declare no competing financial interests. Correspondence and requests for materials should be addressed to Xin Li.

---

**Supplementary Figure 1.**
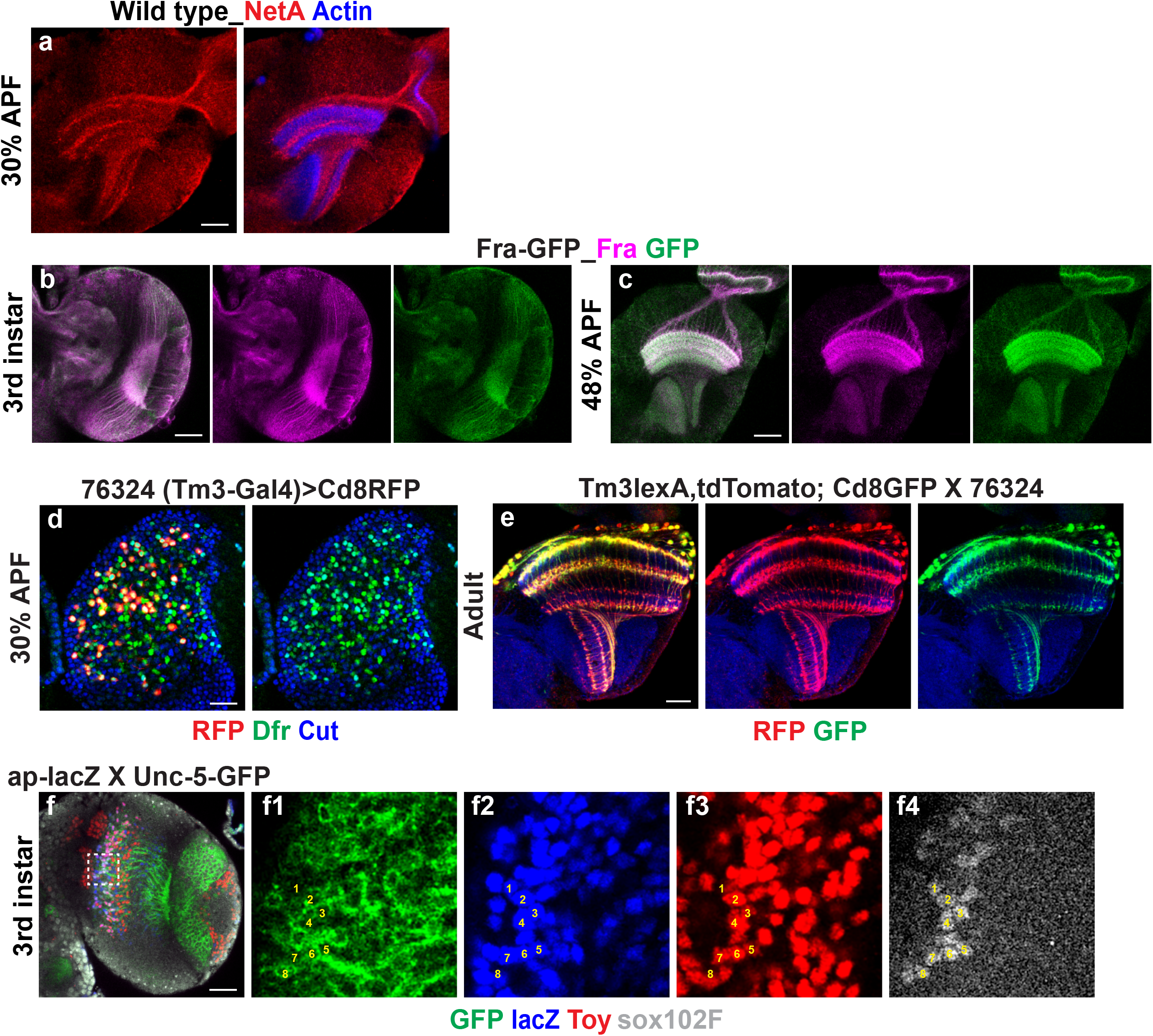
The expression patterns of NetA, Fra, and Unc-5. (a) NetA (red) expression at 30% APF. (b-c) GFP tagged Fra reflects the same expression pattern as stained by anti-Fra. Fra::GFP (green) stained with anti-Fra (purple) at 3^rd^ (b) and 48% APF (c). (d) Characterization of *Tm3-gal4* (BL76324) with known Tm3 molecular markers Dfr (green) and Cut (blue) at 30% APF and known *Tm3-lexA* at adult stage. (f) Tm5a/b (Toy+ Ap+, Sox102F+) neurons do not express Unc-5. Outlined area is shown at higher magnification in (f1)-(f4). Scale bar: 20μm.

**Supplementary Figure 2.**
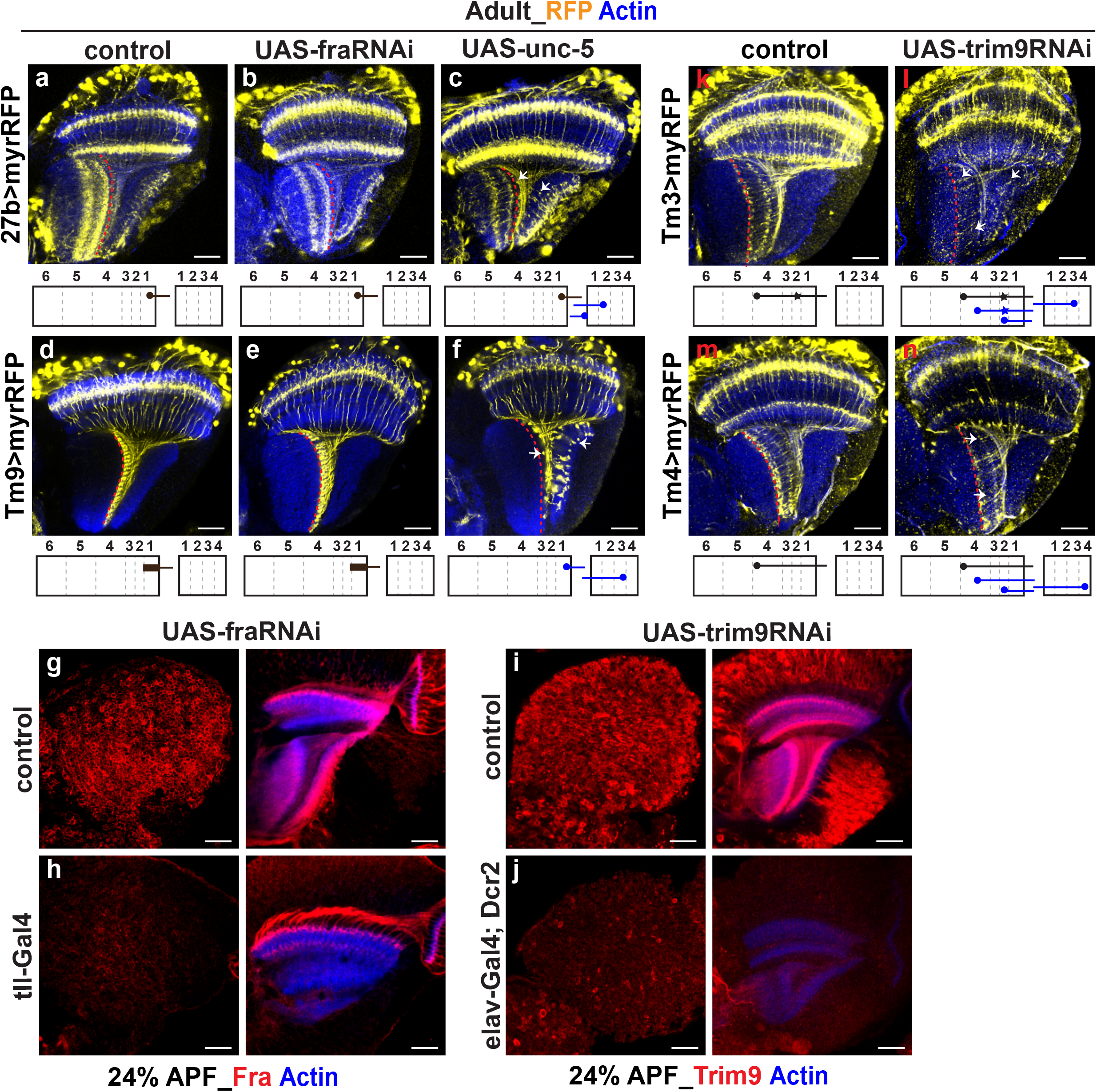
Targeting of Tm neurons when Fra or Trim9 is knocked down or when Unc-5 is mis-expressed. (a-c) Tm1 are labeled by *27b-gal4* with *uas-myrRFP*. The red dashed line labels the wild-type Tm1 axon targeting layer. Arrows point to the mistargeted axon terminals. Schematics are shown below. (a)Wild-type Tm1 (yellow) terminate at distal half of lo1(red dashed line). Note: 27b-gal4 also label other neuron types. (b) Knocking down Fra specially in Tm1 does not affect their targeting. (c) Overexpressing Unc-5 in Tm1. Most Tm1 are repelled from lo1. (d) Wild-type Tm9 terminate at lo1 (red dashed line) with the cylinder-like shape. (e) Knocking down Fra specifically in Tm9 does not affect their targeting. (f) Overexpressing Unc-5 in Tm9. Tm9 are repelled from lo1. (g-h) Knocking down efficiency of *UAS-fraRNAi* is tested. Fra signals are almost all gone in *tll>fraRNAi*. (i-j) Knocking down efficiency of *UAS-trim9RNAi* is tested. Trim9 signals are greatly reduced when knocking down in all neurons. (k-l) Knocking down Trim9 specifically in Tm3 results in mistargeted Tm3. (m-n) Knocking down Trim9 specifically in Tm4 results in mistargeted Tm4. Therefore, Trim9 is also required for Tm3 and Tm4 targeting. Scale bar: 20μm.

**Supplementary Figure 3.**
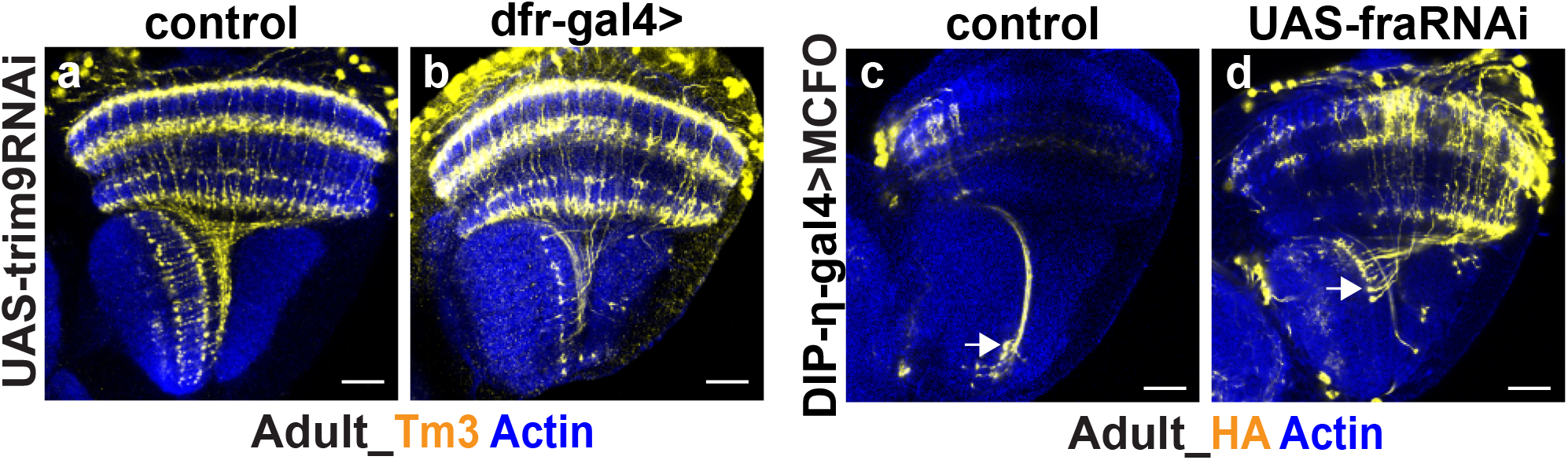
Trim9 is required for Tm3 axon targeting. (a) Wild-type Tm3. (b) Knocking down Trim9 by *dfr-gal4* results in similar defects as knocking down Fra, consistent to the fact that Trim9 is the downstream of Fra. (c-d) MCFO to label *DIP-η-gal4*-expressing cells. (c) Tm2 terminal at lo2 is marked by white arrow. (d) Tm2 targeting is not affected without fra. Scale bar: 20μm.

**Supplementary Figure 4.**
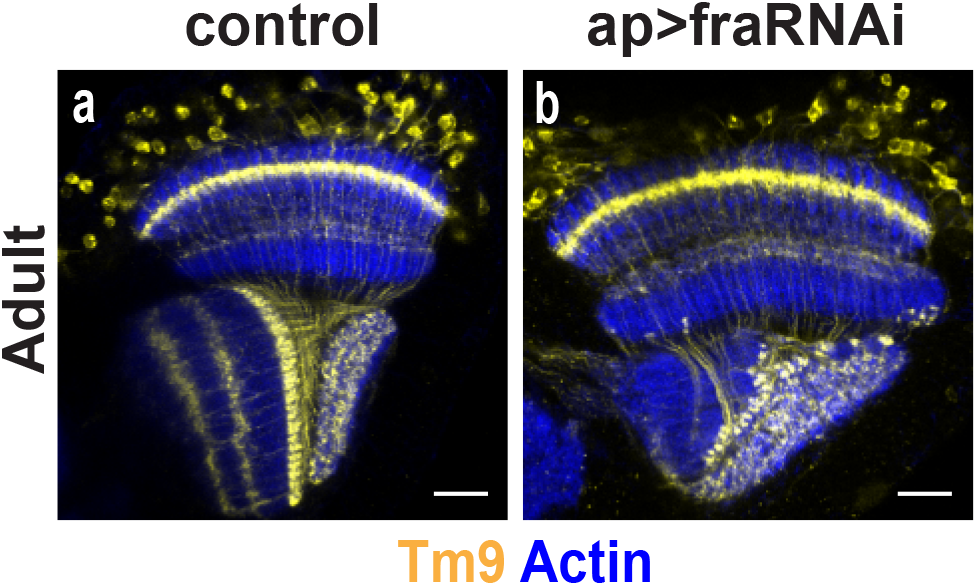
IOC is disrupted when Fra is knocked down in all Ap expressing neurons. (a-b) Tm9 are labeled by *Tm9lexA>lexop-tdTomato*. Note: this driver also labels other neuron types. Tm9 terminate at lo1 and show cylinder-shaped axon terminal. (a) Wild-type Tm9. (b) Knocking down Fra by *ap-gal4* results in disrupted IOC. Scale bar: 20μm.

**Supplementary Figure 5.**
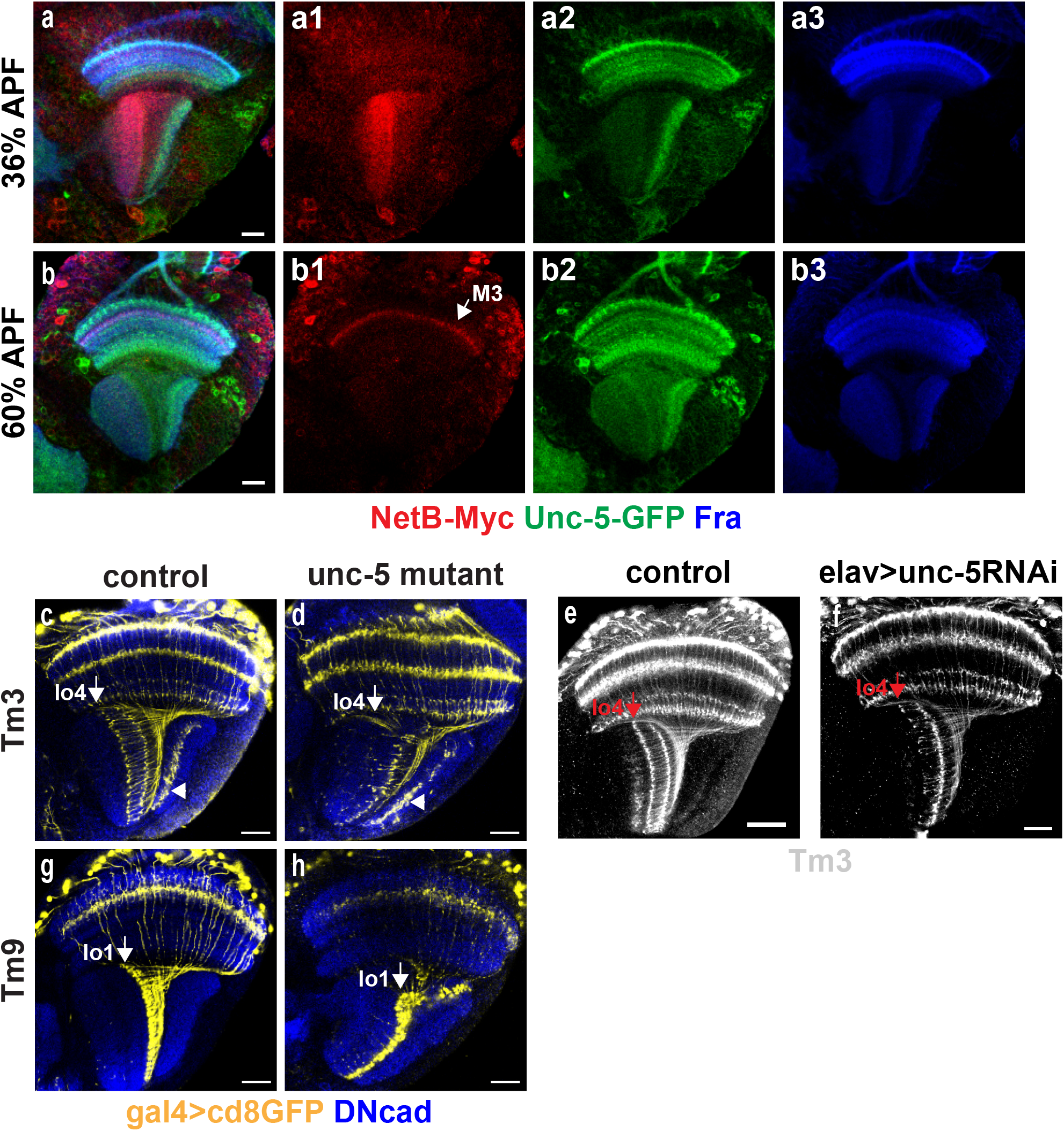
Expression pattern of Unc-5 and Tm targeting defects with loss of unc-5. (a) Unc-5 (green), Fra(blue) and NetB (red) are co-stained at 36% APF. (b) Unc-5 (green), Fra(blue) and NetB (red) are co-stained at 60% APF. Lobula NetB are gone in lobula at this stage. (c-d) Tm3 are labeled by *Tm3-Gal4>cd8GFP*. Note: this driver also labels non-Tm3 neurons that terminate at lop (arrowhead). (c) Tm3 in wild-type. (d) Tm3 targeting in *unc-5* mutants. IOC is disrupted, and Tm3 are mistargeted. (e-f) Tm3 are labeled by *Tm3-lexA>lexop-tdTomato*. (e) Tm3 in wild-type. (f) Knocking down Unc-5 in all neurons results in mistargeting of Tm3. (g) Tm9 in wild-type. (h) Tm9 targeting in *unc-5* mutants. IOC is disrupted, Tm9 still target at lo1, but the distal part of lo1 is mis-located to the surface of lop. Scale bar: 20μm.

**Supplementary Figure 6.**
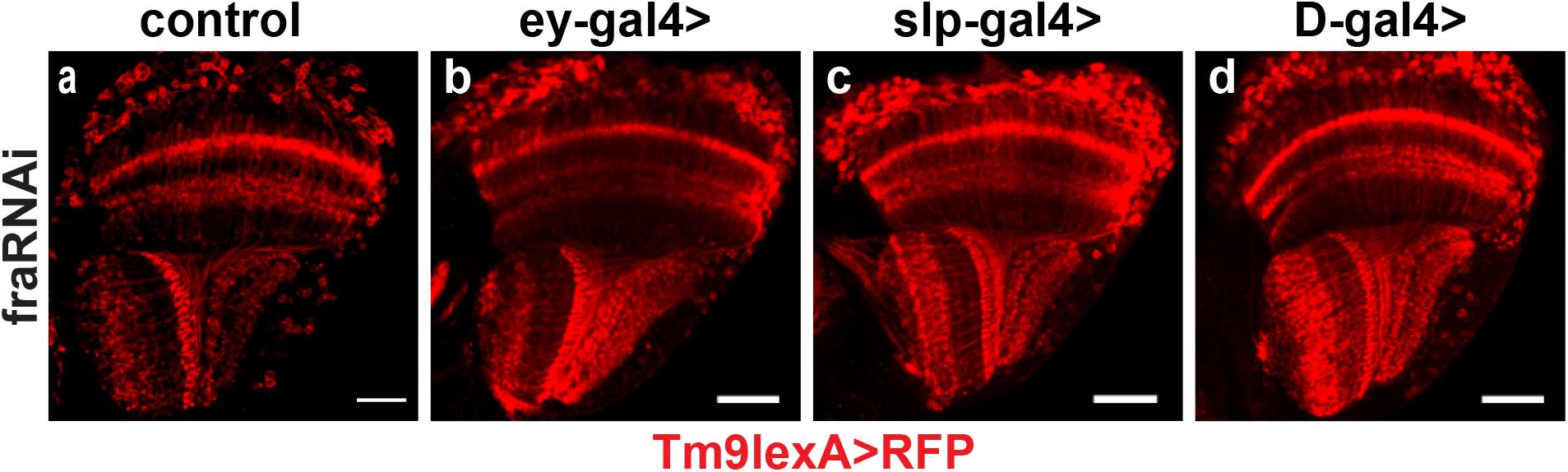
Fra expressed in neurons born in Ey and later stages is not required for IOC formation. (a-d) Tm9 are labeled by *Tm9lexA>lexop-tdTomato* (red). (a) Tm9 in wild-type. (b-d) Tm9 marked IOC is not affected when knocking down fra in neurons born at ey, slp, D temporal windows. Scale bar: 20μm.

**Supplementary Figure 7.**
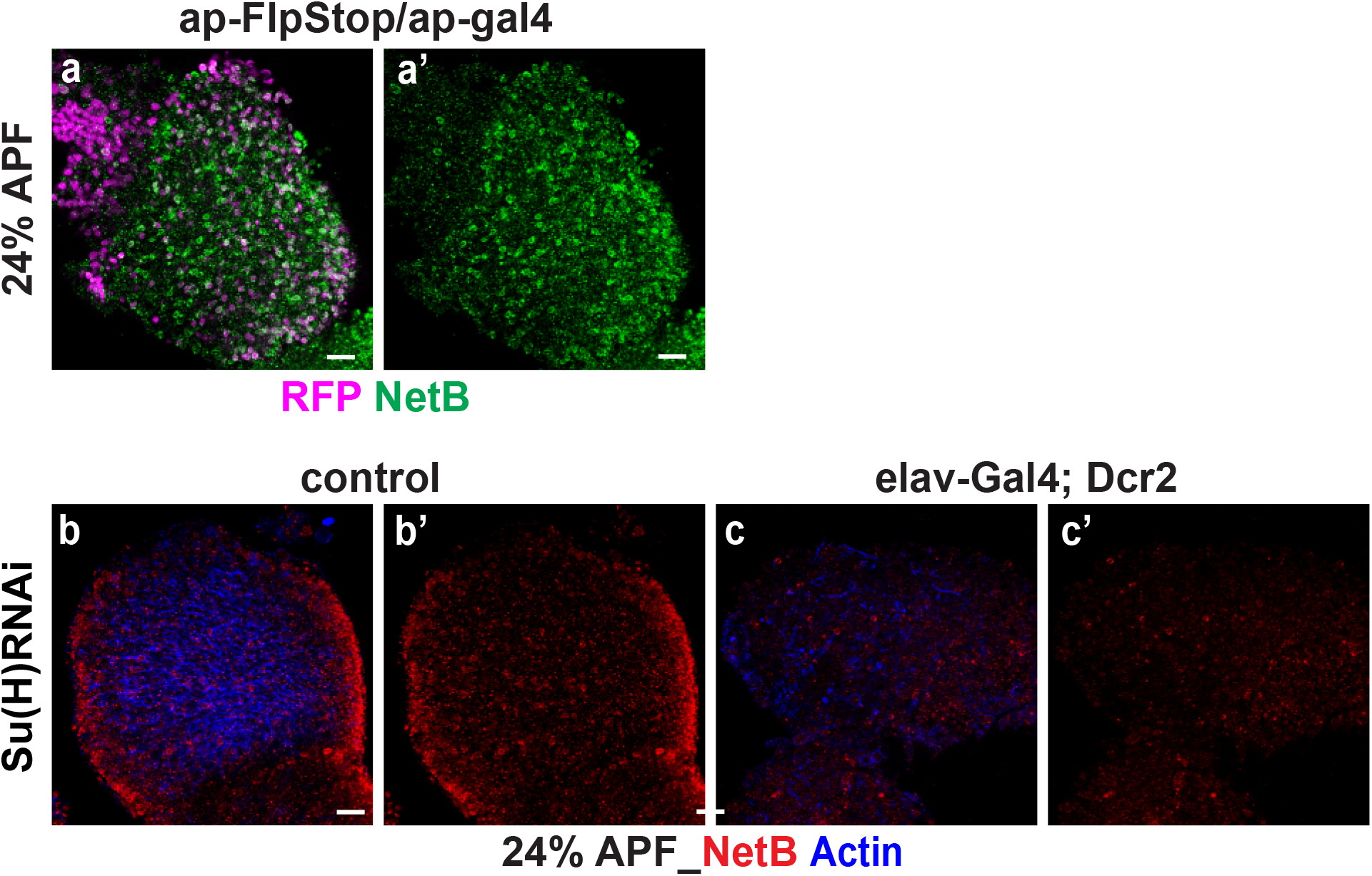
Su(H) but not Ap is required for NetB expression. (a-a’) ap mutants clones are generated by FlpStop. RFP (purple) mark the mutant clones. Some of them are colocalized with NetB signal, indicating that ap is not required for NetB expression. (b-c) Knocking down Su(H) by RNAi abolishes NetB signal at medulla cortex. Optic lobe structure is severely damaged in *elav-gal4;dcr-2>Su(H) RNAi*. Scale bar: 20μm.

